# Diverse Intestinal Injuries Drive Heterogeneous Transcriptional Responses and Limited Reactivation of Developmental Gene Programs in Human Enteroids

**DOI:** 10.64898/2026.08.21.746334

**Authors:** Jonathan W. Villanueva, Yu-Hwai Tsai, Angeline Wu, Carson Caldwell, Abigail Vallie, Mira Buerk, Sha Huang, Jason R. Spence

## Abstract

The murine intestine reactivates developmental gene programs following various forms of damage *in vivo* and *in vitro*; however, injury response mechanisms used by the human intestine remain unclear. Using adult human small intestinal epithelium-only organoids (‘enteroids’), we characterized the early response to eight injury conditions and injury-associated signaling pathways (P53, PGE2, YAP, TGFβ) to interrogate whether human developmental genes were activated. P53 activation and decreased proliferation were common features across treatments. Most (7/8) injuries did not activate human development genes. Butyrate is a notable exception given it inhibited P53 and promoted a human developmental transcriptional signature. We observe that P53 induces a human adult gene signature while TGFβ and YAP promote a developmental signature. Together our data characterizes various transcriptional responses to injury, supports injury-associated signaling pathways as regulators of human adult and developmental genes, and highlights how our data can be mined to predict injury-specific interventions for epithelial protection.

## Introduction

The small intestinal epithelium is single layer of cells that lines the inside of the intestine and forms a complex architecture organized into functional units called crypts and villi. Crypts harbor the intestinal stem cells (ISCs) that are critical for maintaining epithelial cell populations, whereas villus structures consist of absorptive and secretory cells that are responsible for metabolic function, nutrient absorption and maintaining tissue integrity. The intestinal epithelium also functions as a protective barrier that shields the host from exposure to microbes, food and harmful stimuli including pathogens^1^, environmental toxins^2,3^, cytokines^4^, and pharmaceuticals^5,6^. To maintain normal cell turnover and tissue integrity following damage, the intestine requires robust mechanisms to facilitate homeostasis and tissue regeneration following injury. Breakdown of these repair programs can inhibit the intestine’s ability to absorb nutrients and ultimately drive intestinal disease^7–9^.

During homeostasis, intestinal stem cells (ISCs) are constantly proliferating to produce progeny that differentiate to maintain secretory and absorptive cell populations^10^. Following damage, ISCs are often lost and differentiated epithelial cells are capable of de-differentiating and adopting stem cell functions to regenerate damaged tissues^11–13^. Recent studies have built upon this idea of injury-induced plasticity by highlighting a process called “fetal-like reversion”, where the intestine reactivates gene programs utilized during development as a part of the regenerative process^14–20^. Fetal-like reversion is characterized by the reduction or loss of adult ISC marker genes (*Lgr5*, *Olfm4, etc.*) and activation of genes enriched in the fetal mouse intestine (*Sca1*, *Tacstd2*, *Clu*, *Spp1*, etc.) during repair of adult intestinal tissue^14^. Many *in vivo* and *in vitro* mouse models^15–18,21^ of intestinal injury (helminth infection^15^, chemical treatment^16,17^, irradiation^15,17^) have been shown to induce a fetal-like gene signature, mechanistically linked to signaling pathways including YAP^16^, TGFβ^17^ and IFNγ^15^. However, it remains unclear whether a fetal-like gene program is also leveraged by the adult human intestine in response to damage, especially given that rodent cells can respond to injury differently than human cells^22–25^.

Here, we sought to interrogate how the intestinal epithelium responds to multiple forms of damage and evaluate how genes utilized in injury-associated signaling pathways and development are leveraged following injury. To do this, we treated human enteroids with eight different stimuli to induce various types and degrees of cell stress/injury: Mycophenolic acid (MA)^5,26^; Oxaliplatin^27,28^; Arsenic^3,29^; Deoxynivalenol (DON)^30,31^; Ibuprofen^32,33^; Ethanol^34,35^; 5-Fluorouracil (5FU)^27,36^; Butyrate^37,38^. We also treated enteroids with modulators for signaling pathways that have been implicated in the injury response (P53^39^, PGE2^40^, YAP^41^, TGFβ^17^). Finally, we leveraged sequencing datasets from the developing and adult human intestinal epithelium to identify a consensus set of 140 genes commonly enriched in the fetal human intestine, both *in vitro* and *in vivo,* to determine if adult human enteroids activate a fetal-like gene program in response to injury. Our results show that P53 activation and reduced proliferation are common features of the enteroid injury response. Furthermore, human adult enteroids do not significantly activate a human fetal gene signature in seven out of the eight treatments tested. Butyrate was a unique exception because it was the only condition to promote a fetal gene signature and suppress P53 activity. Interrogation of signaling pathways revealed that TGFβ and YAP activation promote a human fetal-like gene signature, whereas P53 activation in human enteroids pushes them towards an adult transcriptional signature. Finally, we interrogated sequencing data for individual injury conditions and found that DON treated enteroids downregulate genes associated with arachidonic acid metabolism. Supplementing cultures with arachidonic acid, or its downstream product PGE2, provides a protective effect against DON-induced injury, while the other forms of injury were not protected. To easily interrogate the 140 bulk RNA-seq datasets created for this study, we also developed a publicly accessible database, called HD-REPAIRD (https://spence-lab.shinyapps.io/HD-REPAIRD/) as a rich resource for the research community to utilize.

Taken together, our findings suggest that human intestinal epithelial cells do not robustly activate fetal gene programs immediately following exposure to most injury stimuli, at least in part, due to the antagonistic relationship between P53 and the human fetal gene program. Beyond establishing this paradigm, our identification of arachidonic acid and PGE2 as injury-specific protective molecules demonstrates how mechanistic insight gained from individual injuries can uncover protective mechanisms that guide the development of novel therapeutic strategies.

## Results

### Human enteroids exhibit unique changes in gene expression following exposure to various injury stimuli

To investigate how human intestinal cells respond to various forms of injury, we treated human adult enteroids with eight different damage/stress stimuli for 48 hours (n= 3 different patient derived lines) (**Figure 1A**). We focused on agents that are known to induce various forms and degrees of cell stress/injury consisting of chemotherapeutics (100 uM oxaliplatin^27,28^, 500 uM 5-Fluorouracil – 5FU^27,36^), anti-inflammatory agents (100 ug/mL mycophenolic acid - MA^5,26^, 400 ug/mL ibuprofen^32,33^), a heavy metal (20 uM arsenic^3,29^), a mycotoxin (40 uM deoxynivalenol – DON^30,31^), a short chain fatty acid (8.73 mM butyrate^37,38^), and 5% ethanol^34,35^ which induces oxidative cell stress. To visualize cells actively undergoing apoptosis, a caspase-3/7 detection reagent was added to the media during the injury period to fluorescently label apoptotic cells. At the end of the 48-hour incubation, brightfield and fluorescent images were taken. To quantify changes in apoptosis and enteroids size, CellPose^42^ was used to define enteroid boundaries and create a mask that is applied to each fluorescent image to calculate the percentage of each enteroid that is positive for fluorescent signal (**Figure 1B-1C; Supplemental Figure 1A-1B**). All injury treatments resulted in a significant decrease in enteroids size (**Figure 1D**), and 6/8 conditions produced an increase in the relative abundance of apoptotic cells (**Figure 1E**). Immunofluorescence (IF) staining for proliferation markers KI67 and CCNB1 showed that 5/8 treatments (oxaliplatin, DON, 5FU, ibuprofen, butyrate) result in a severe decrease in proliferation (**Figure 1F; Supplemental Figure 1C**). Taken together, these results support that the eight different types of injury/stress impact enteroid size, proliferation and apoptosis to varying degrees.

**Figure 1:**
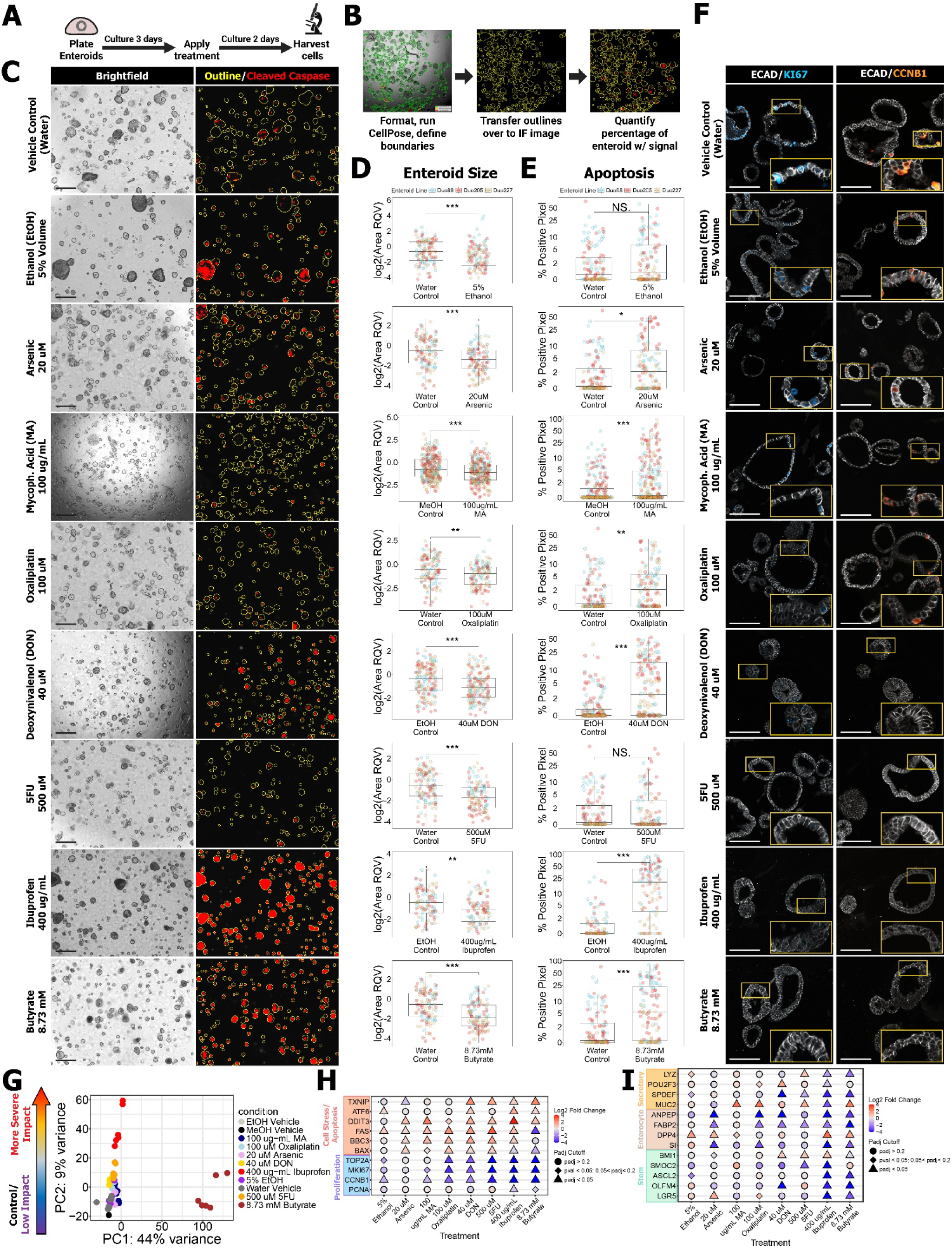
Human adult enteroids exhibit injury-specific changes in proliferation and cell death. (**A**) Diagram of injury treatment timeline (Created with BioRender.com). (**B**) Schematic outlining the image analysis pipeline for CellEvent Caspase 3/7 assay. (**C**) Brightfield (left) and analyzed fluorescent images (right) highlighting enteroid boundaries (yellow) and pixels positive for fluorescent signal (red). Scale bar = 500 um. (**D,E**) Quantification of changes in enteroid size relative to control (D; RQV = relative quantitative value) and percentage of each enteroid that is positive for CellEvent Caspase 3/7 reagent signal (E). Enteroid line denoted by color. Significance determined using two-sided Wilcoxon test. *p < 0.05, **p < 0.01, ***p < 0.001 (**F**) IF images for proliferation markers KI67 (Blue) and CCNB1 (Orange). Cell boundaries visualized using ECAD (white). Scale bar = 100 um. (**G**) PCA generated using gene expression profiles from enteroids treated with eight injury stimuli and three vehicle controls. (**H, I**) Dot plots visualizing changes in expression of gene markers for proliferation (H), apoptosis/cell stress (H), and different intestinal cell populations (I). Datapoints formatted according to DESeq2 results where color represents the log2 fold change and shape signifies statistical significance. All experiments were carried out in n=3 independent biological replicates, denoted as Duo88, Duo205 and Duo 227.

To characterize injury-induced changes in gene expression, we harvested cells for RNA isolation and performed bulk RNA-seq. Principal component analysis (PCA) revealed that the largest percentage of the gene variance (PC1) is driven by gene expression changes induced by butyrate treatment (**Figure 1G, Supplemental Figure 1D**). Samples next sorted along PC2 in a gradient of injury severity, ranging from vehicle controls to more severe impact treatments like 5FU and ibuprofen. Differential expression analyses were performed using DESeq2^43^ and used to interrogate changes in the expression of genes related to cell stress, cell death, and proliferation. Our analysis revealed that enteroids exhibited variable activation of cell stress (*TXNIP^44^*, *ATF6^45^*, *DDIT3^46^*) and cell death (*FAS^47^*, *BBC3^48^*, *BAX^49^*) genes and decrease in proliferation genes (*TOP2A^50^*, *MKI67^51^*, *CCNB1^52^*, *PCNA^51^*; **Figure 1H**). Interestingly, we found that while butyrate treatment induced a strong increase in cell stress markers, it also decreased the expression of apoptotic markers. This is in line with the mixed literature on butyrate where it has been cited as having a protective effect in some contexts^37,53^, while being harmful in other contexts^37,54^.

Analysis of cell-specific markers revealed a robust loss in ISC markers (*SMOC2^51^*, *ASCL2^51^*, *OLFM4*^51^, *LGR5*^51^) and secretory markers (*LYZ^55^*, *POU2F3* ^51^, *SPDEF*^51^) following ibuprofen and butyrate treatment (**Figure 1I**). Most injury treatments (5/8 treatments) decreased the expression of enterocyte markers (*SI^56^*, *FABP2*^51^, *ANPEP*^51^). Interestingly, the enterocyte marker *DPP4^57^* was elevated in 4/8 treatment conditions (**Figure 1I**), suggesting a unique role for *DPP4* during the injury response. We also observed an elevation in the goblet cell marker *MUC2*^51^ (4/8 treatments) in multiple conditions (**Figure 1I**). Some treatments induced unique injury-specific elevations for certain markers. Notably this includes arsenic treatment which elevates *LGR5*, DON treatment which elevates *POU2F3*, and 5FU treatment that promotes *LYZ* expression (**Figure 1I**). Taken together, these results highlight that different cell stressors lead to unique gene expression responses.

### Human enteroids exhibit injury-specific gene expression changes that converge on reduced proliferation

To determine if there were common pathways activated across injuries, we first performed a series of differential expression analyses comparing each injury condition to the appropriate vehicle control (**Figure 2A**). From this, we identified 251 genes that were upregulated in response to four or more of the eight injury conditions (log2 fold change > 1, padj<0.05, baseMean > 10; **Figure 2B, Supplemental Table 1**). A KEGG pathway analysis using Enrichr^58–60^ revealed a statistical enrichment for genes relating to signaling pathways affiliated with the injury response (Hippo, P53, MAPK), which suggests that these pathways are commonly utilized across injury types and/or severities (**Figure 2C**).

**Figure 2:**
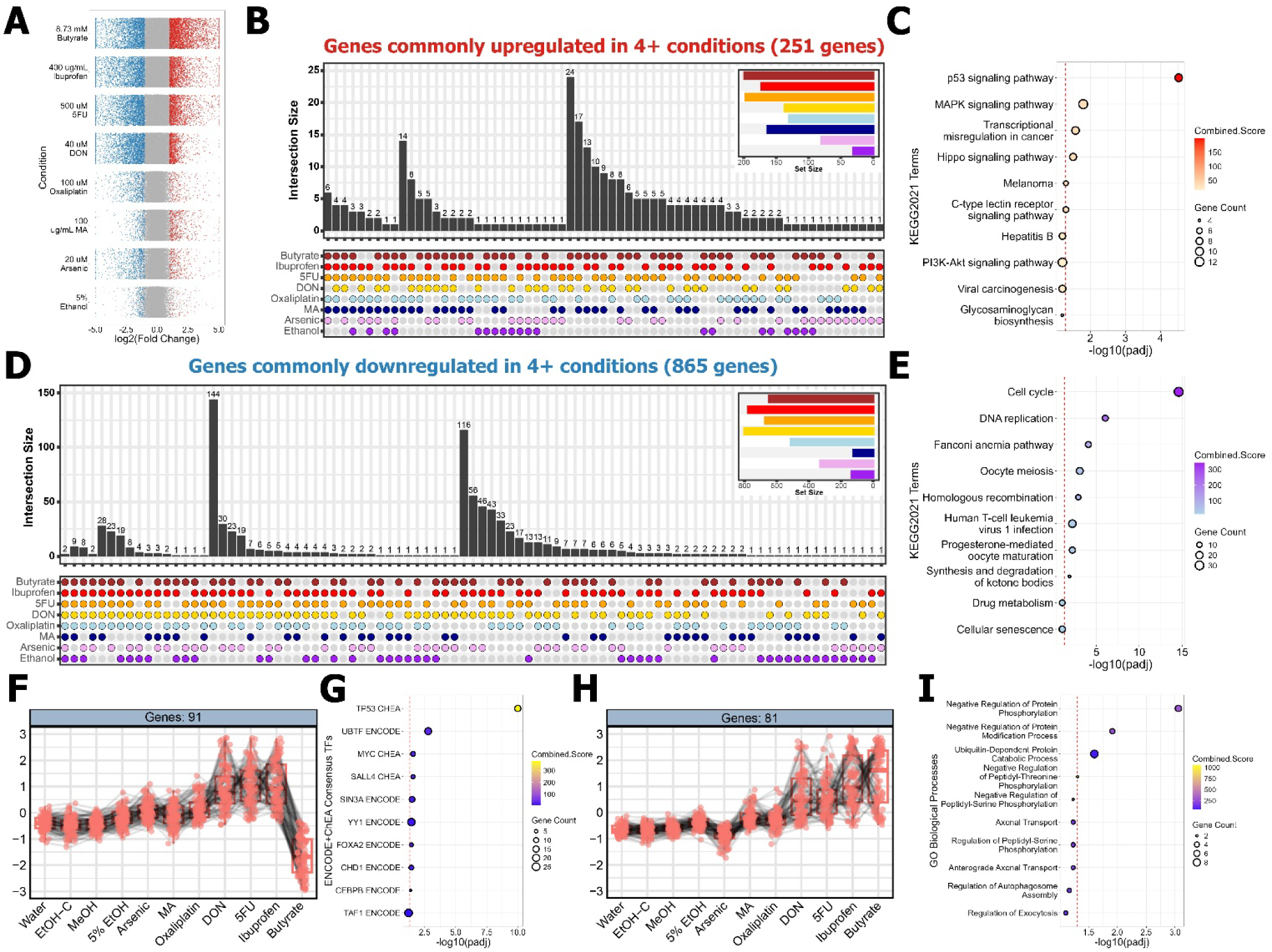
Commonly differentially expressed genes are enriched for genes affiliated with P53 signaling and proliferation. (**A**) Volcano plots visualizing up- (red) and downregulated (blue) genes for each injury treatment. (**B, D**) Upset plots visualizing the overlapping intersections of genes upregulated (B) or downregulated (D) in four or more conditions. Set size (top right corner) represents the total number of genes from each condition that are represented in the plot. (**C, E**) Dotplots visualizing results from Enrichr KEGG pathway analyses performed using commonly up- (C) or downregulated (E) genes. Dotted red line illustrates padj = 0.05. Dot size represents the number of genes from the query lists assigned to each KEGG term. Dot color represents the combined score calculated by Enrichr as a metric of significance. (**F, H**) Z-score values of gene expression for 2/30 gene modules defined by DEGReport. Gene modules group genes together with similar patterns of expression across control (Water, EtOH-C, MeOH) and injury (5% EtOH, Arsenic, MA, Oxaliplatin, DON, 5FU, Ibuprofen, Butyrate) conditions. Each datapoint represents a gene where the black lines connect datapoints for the same genes across all conditions. (**G, H**) Dotplots visualizing results from Enrichr gene enrichment analyses using ENCODE ChEA Consensus Transcription Factor (G) and GO Biological Processes (I) terms.

In a similar analysis, we identified 865 genes that were commonly downregulated in response to at least four of the injury stimuli (log2 fold change < -1, padj<0.05, baseMean > 10; **Figure 2D, Supplemental Table 2**). From this list, *TRIM17* and *POGLUT3* were the only genes downregulated in all eight treatments, suggesting that their functional roles are broadly applicable across injuries. Results from a KEGG pathway analysis highlight that the 865 commonly downregulated genes contain an enrichment for genes related to DNA replication and the cell cycle (**Figure 2E**). This result is concordant with the enrichment of P53 signaling genes in the commonly upregulated genes list (**Figure 2C**) given P53 is a well-known negative regulator of cell cycle progression^61^.

Next, we sought to define groups of genes that exhibit similar patterns of expression across injury treatments. To do this, we used DESeq2 to perform a likelihood ratio test and filtered for genes that exhibited significant variation (padj<0.05) across conditions. We next sorted genes in ascending order based on adjusted p-values and subset for the top 3000 genes. Using the R package DEGReport^62^ 30 modules were identified, each consisting of >50 genes, that depicted 30 different patterns of gene expression across injury treatments (**Supplemental Figure 2, Supplemental Table 3**). Throughout gene modules, we observed that the data variation is primarily driven by gene expression differences in samples treated with DON, 5FU, ibuprofen, butyrate (**Supplemental Figure 2**). For example, one group is composed of genes that are most strongly elevated in response to DON, 5FU and ibuprofen treatment, but strongly downregulated following butyrate treatment (**Figure 2F**). A gene enrichment analysis using ENCODE and ChEA consensus transcription factor terms found that this group contained a significant enrichment for genes with P53 binding sites (**Figure 2G**). This is concordant with our finding from **Figure 2C**. The decreased expression of P53 target genes following butyrate treatment is also consistent with literature showing that butyrate suppresses P53 actvitity^63^. A separate group of genes consists of transcripts that are most highly upregulated following DON, 5FU, ibuprofen, and butyrate treatment (**Figure 2H**). A GO analysis revealed enrichment for regulators of protein modifications/phosphorylation, suggesting injury-specific alterations at the protein level (**Figure 2I**). These two examples highlight the valuable insight that our dataset reveals about the shared and injury-specific impacts of various injury treatments on human intestinal biology.

### P53 signaling is commonly activated in human enteroids exposed to various stress/injury stimuli

Given our results highlighting Hippo/YAP and P53 associated genes during the injury response (**Figure 2**), and the well-established role for these and other pathways during injury-repair (P53^39,64^, YAP^16,65^, TGFβ^17,66^, PGE2^40,67^), we leveraged agonists to interrogate how specific activation of these signaling pathways compared to the gene signatures of human enteroids exposed to injury (**Figure 2**). Enteroids were treated with a LATS inhibitor (LATSi – promotes nuclear localization of YAP), Nutlin (inhibits MDM2-induced degradation of P53), TGFβ and PGE2 for 48 hours using the same experimental paradigm described for injury stimuli (**Figure 1A**). RNA was isolated from cells for bulk RNA-seq analysis (**Figure 3A, Supplemental Figure 3A**) where we utilized DESeq2 to identify upregulated and downregulated genes in response to targeted signaling pathways (**Figure 3B**). We next leveraged the Broad Institute Gene Set Enrichment Analysis (GSEA)^68,69^ tool to evaluate how genes activated by different signaling pathways change in response to our eight injury stimuli. Normalized enrichment scores (NES) from this analysis are visualized using a radar plot where each node indicates whether a given treatment activates a specific set of signaling pathway genes (**Figure 3C** – Black node signifies no significant enrichment, Red node signifies positive enrichment, Blue node signifies negative enrichment). Notably, we found that genes stimulated by P53 were positively enriched in 7/8 injury conditions, with butyrate being the only exception (**Figure 3C**). This finding corresponds to the gene module analysis (**Figure 2F-2G**), where P53 associated genes were downregulated in the butyrate condition. Genes activated by TGFβ were positively enriched in 4/8 conditions whereas genes stimulated by LATSi-induced activation of YAP were only positively enriched in 2/8 conditions. PGE2 stimulated the smallest gene expression changes (**Figure 3B**), and the PGE2 gene signature was positively enriched in the MA treatment group, making MA the only stimulus that activated genes downstream of all four signaling pathways. Of the 251 genes commonly upregulated across injuries (**Figure 2B**), 87/251 (34.7%) were significantly elevated (log2 fold change > 1, padj<0.05) by Nutlin treatment (**Figure 3D**), while 61/251 (24.3%) were significantly upregulated (log2 fold change > 1, padj<0.05) by TGFβ treatment (**Figure 3D**). These results support that P53 signaling is a major driver of the injury-response signatures observed in the injury conditions.

**Figure 3:**
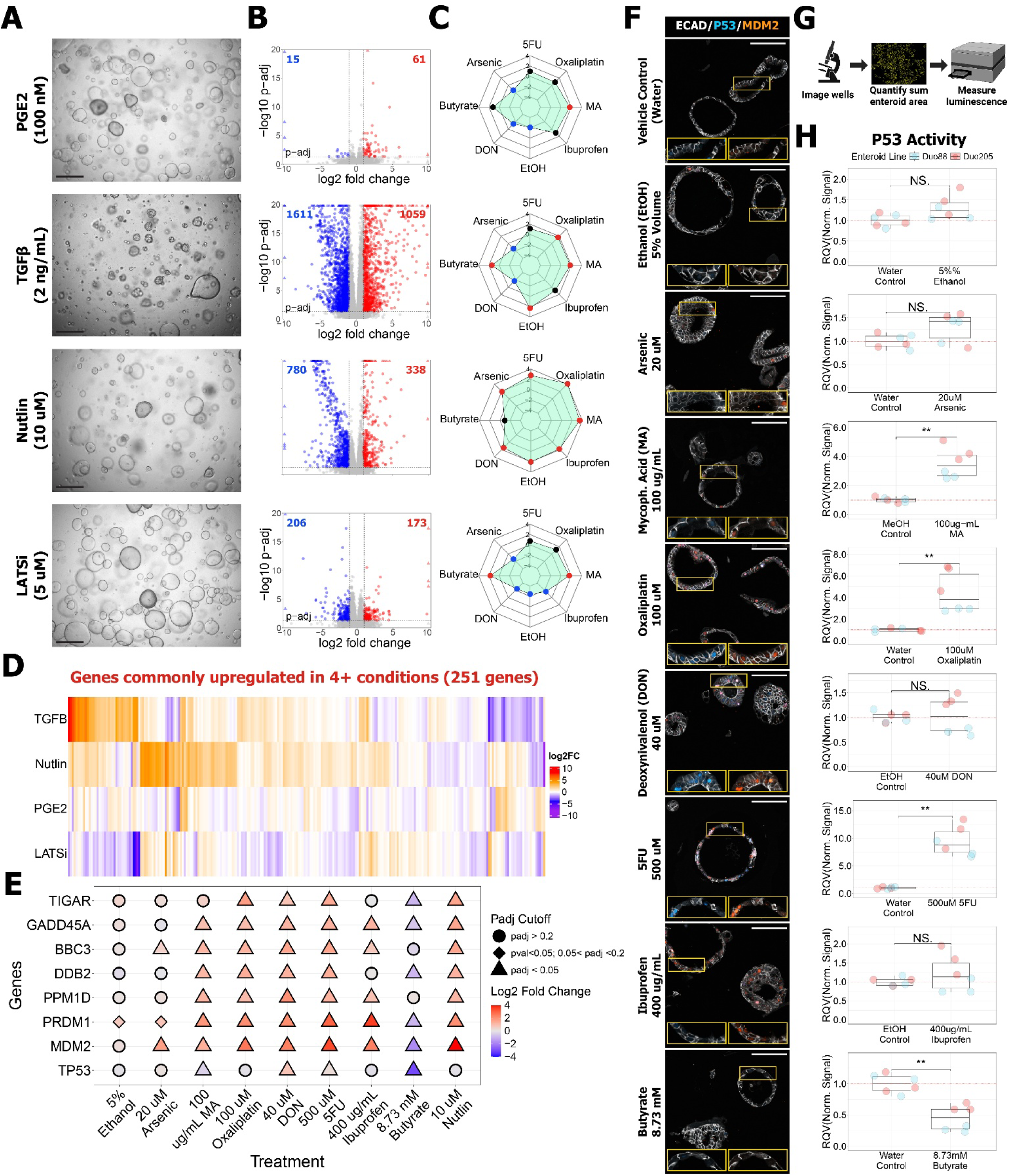
P53 signaling is activated to various degrees across most injury treatments. (**A**) Brightfield images of enteroids treated with PGE2, TGFβ, Nutlin and a LATS inhibitor (LATSi). Scale bar = 500 um. (**B**) Volcano plots visualizing up- (red) and downregulated (blue) genes for each treatment relative to vehicle control. Black dotted line represents padj = 0.05. (**C**) Radar plots displaying normalized enrichment scores (NES) calculated using the Broad Institute GSEA tool to evaluate activation of genes associated with injury-associated pathways. Red = significant positive enrichment. Blue = significant negative enrichment. Black = no enrichment. Statistical significance = FDR q-value < 0.05 (**D**) Heatmap visualizing how activation of the four signaling pathways affects the expression of genes commonly elevated by injury conditions established in Figure 2B. Color scale represents the log2 fold change calculated by DESeq2. (**E**) Dot plot visualizing differential expression results (DESeq2) for P53 response genes in enteroids treated with eight injury stimuli and Nutlin. Datapoint color represents the log2 fold change and shape indicates statistical significance. (**F**) IF images for P53 (Blue), MDM2 (Orange), and ECAD (white). Scale bar = 100 um. (**G**) Diagram outlining the strategy for quantification of P53 activity using P53 luciferase reporter. (**H**). Quantification of relative luciferase activity normalized to enteroid input. Enteroid line denoted by color. Significance determined using two-sided Wilcoxon test. All experiments were carried out in n=3 independent biological replicates, denoted as Duo88, Duo205 and Duo 227, except for Figure 3H with n=2 independent biological replicates. *p < 0.05, **p < 0.01, ***p < 0.001.

We also directly interrogated expression of *TP53* and several downstream target genes (**Figure 3E**). While *TP53* target genes are prominently activated across most injury conditions, we only observed a significant increase in *TP53* expression following DON and 5FU treatment (**Figure 3E**). Ethanol and Arsenic treatment showed the weakest activation of P53 target genes amongst all the conditions (**Figure 3E**). In order to further interrogate P53 activation in stress/injury conditions, we carried out IF staining for P53 and MDM2, a well-described transcriptional target of P53^70,71^. We observed very little nuclear staining for either protein in the control condition, and staining was absent in the butyrate conditions. However, obvious nuclear staining was present in all other stress/injury conditions. The most intense staining was observed in enteroids treated with oxaliplatin, 5FU and DON (**Figure 3F, Supplemental Figure 3B**).

Finally, to directly measure P53 transcription factor activity in response to each of the eight injury stimuli, we generated P53 luciferase reporter enteroid lines (**Supplemental Figure 3C**). Prior to performing the luciferase assay, brightfield images were taken of each well and used to normalize the luminescent signal relative to the cellular input (**Figure 3G**). Results show that 5% EtOH (p-val = 0.093) and arsenic (p-val = 0.13) treatment produced a minor, but not statistically significant increase in P53 activity (**Figure 3H**). MA, oxaliplatin and 5FU produced robust increases in P53 activity while butyrate significantly decreased P53-induced luciferase signal (**Figure 3H**). Interestingly DON and Ibuprofen did not produce a robust change in luciferase signal (**Figure 3H**). To evaluate whether the luciferase transcript was produced under these conditions, we leveraged RT-qPCR to confirm that DON^72^, but not ibuprofen, significantly increased the production of luciferase mRNA (**Supplemental Figure 3D**). This result suggests that DON also increases P53 activity but that its role as a ribotoxic agent^72^ may impair the production of luminescent signal. Regarding ibuprofen, prior literature has shown that P53 and ATF4 bind and activate a similar set of genes^73^. Characterization of P53-specific, ATF4-specific and shared gene targets revealed that ibuprofen strongly upregulates ATF4-specific and shared genes but only upregulates a minority of P53-specific genes (**Supplemental Figure 3E**). This result supports a model whereby human enteroids exposed to ibuprofen likely respond to injury by activating ATF4 to induce a similar, but distinct, transcriptional program to P53. Together, our data shows that alteration of P53 signaling is a common response across a wide range of stress/injuries, except for butyrate and ibuprofen treatment.

### Defining consensus human fetal and adult gene lists using human datasets derived from in vitro and in vivo sources

In mice, *in vivo* injury has been shown to activate a YAP-dependent repair process, where the intestinal epithelium upregulates genes that are enriched in the fetal mouse intestine, a process called fetal-like reversion^16,74^. Therefore, we sought to evaluate whether adult human enteroids activate human fetal genes following injury. To do this, we first aimed to define a “human fetal intestinal gene signature” as published fetal gene signatures are derived from mouse models^16,75^ which have developmental differences from humans^76–78^. We utilized a two-pronged approach to define fetal human intestinal genes that are present both *in vitro* and *in vivo.* First, we performed bulk RNA-seq on four human adult enteroid lines and six human fetal enteroid lines that were established and maintained in identical growth conditions (**Figure 4A, Supplemental Figure 4A**). From this analysis, we identified 761 genes enriched in fetal enteroids (log2 fold change >1, padj<0.05, baseMean > 10; **Supplemental Table 4**) and 1002 genes enriched in adult enteroids (log2 fold change < -1, padj<0.05, baseMean > 10; **Supplemental Table 5**) (**Figure 4B**). Surprisingly, when using GSEA to evaluate how previously published mouse fetal genes^16^ differ in our human fetal enteroids, we found that human fetal enteroids exhibit a significant negative enrichment for mouse fetal genes (**Figure 4C**). This result suggests that the human and mouse fetal gene signatures differ significantly.

**Figure 4:**
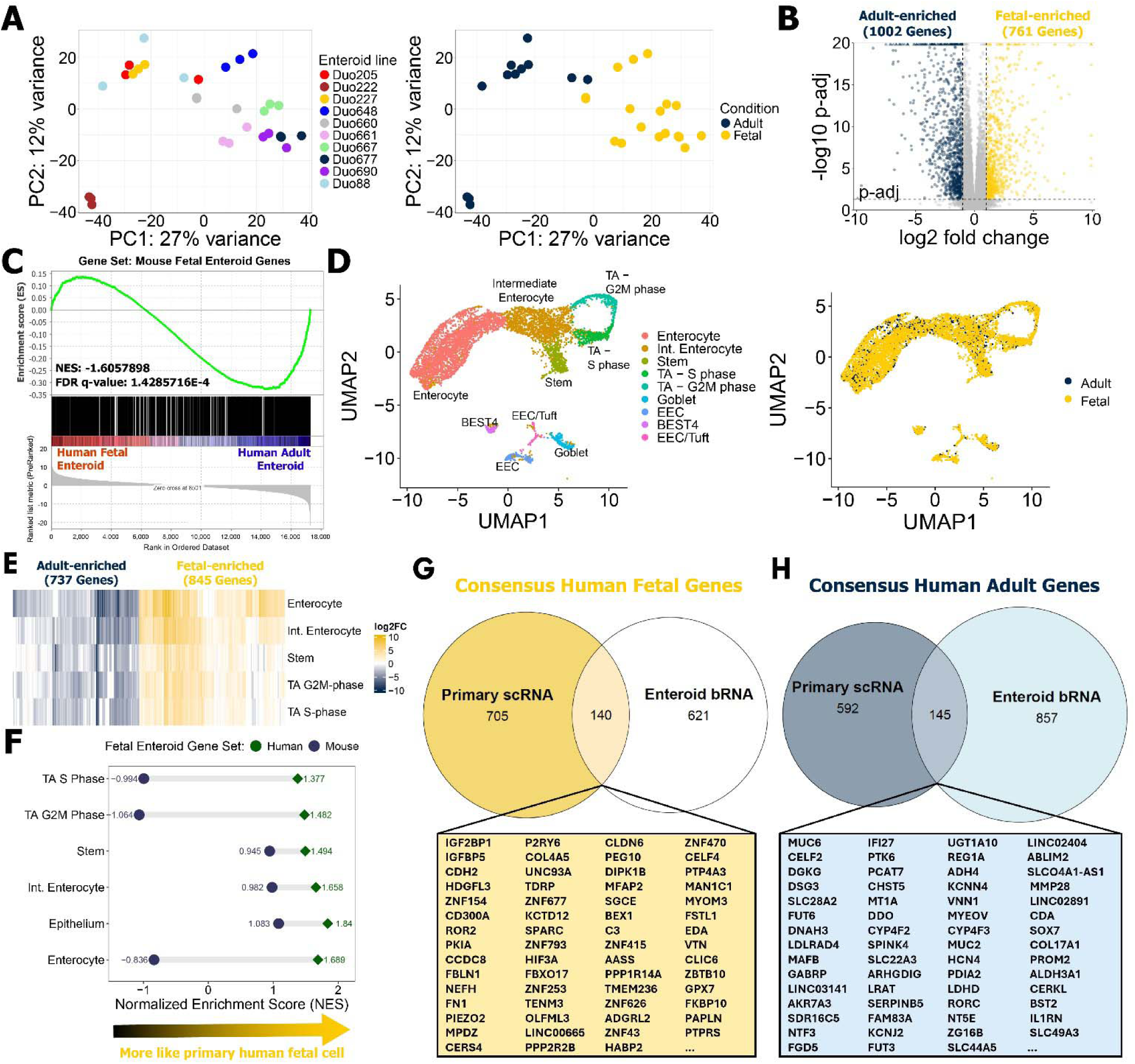
Defining genes enriched in human fetal and adult intestinal cells. (**A**) PCA generated using gene expression profiles from human fetal (6 lines) and adult (4 lines) enteroids at baseline (i.e. no treatment). (**B**) Volcano plot visualizing fetal (yellow) and adult (navy) enriched genes. Black dotted line represents padj = 0.05. (**C**) GSEA enrichment plot evaluating enrichment for mouse fetal enteroid genes in human fetal relative to adult enteroids. (**D**) Uniform manifold approximation and projection (UMAP) of single-cell data previously generated from human primary adult (n=2) and developing (n=7) intestine. (**E**) Heatmap visualizing differences in expression of genes enriched in the human fetal and adult intestine. Color scale represents the log2 fold change calculated by Seurat. (**F**) Dumbbell plot visualizing results from GSEA analysis evaluating enrichment of mouse fetal enteroid genes (dark blue) or human fetal enteroid genes (green) in different cell populations in the primary fetal intestinal epithelium. Shape indicates whether result was statistically significant (FDR q-value < 0.05; diamond) or not (FDR q-value > 0.05; circle). (**G, H**) Venn diagrams highlighting the overlap of fetal (G) and adult (H) genes that were identified using the human enteroid bulk RNA-seq and primary tissue single-cell RNA-seq datasets.

To further refine our gene signature and ensure that it is relevant to the *in vivo* setting as well, we next leveraged previously published single-cell RNA-seq data from human fetal (n=7) and adult (n=2) small intestine^56,57^ (**Figure 4D**, **Supplemental Figure 4B**). We performed a series of differential expression analyses comparing fetal and adult enterocytes, transit amplifying (TA) cells, and stem cells. We focused on these cell populations as these are the primary cell types in the enteroid model. Genes significantly elevated in these fetal cell types (log2 fold change > 1, padj<0.05) were pooled together and further filtered for genes that were elevated in the total fetal epithelium (log2 fold change > 0). We next filtered for genes that were expressed in <5% of adult epithelial cells to ensure they were primarily expressed in a developmental context. This analysis left us with 845 genes (**Supplemental Table 6**) that were enriched in fetal epithelial cell types relative to adult while the inverse analysis defined 737 adult-enriched genes (**Supplemental Table 7**) (**Figure 4E**). Leveraging the GSEA tool, we compared gene expression profiles of primary human fetal epithelial cells (**Figure 4E**) to *in vitro* human fetal (**Figure 4B**) and published mouse fetal enteroid gene lists^16^. NES values are visualized as a dumbbell plot to visualize whether human or mouse fetal enteroid genes are more strongly associated with human fetal epithelial cells derived from primary tissue. Interestingly, we found that there is no positive enrichment of fetal mouse enteroid genes in primary human fetal epithelial cells, but there is a positive enrichment for human fetal enteroid genes (**Figure 4F**). This data supports that there are significant differences between mouse and human regarding what genes are enriched in the fetal epithelium relative to adult. Comparing the fetal/adult gene lists from the enteroids (**Figure 4B**) and primary tissue (**Figure 4E**) analyses, we identified 140 consensus human fetal intestine genes (**Supplemental Table 8**) and 145 consensus human adult intestine genes (**Supplemental Table 9**) that were commonly enriched *in vitro* and *in vivo* (**Figure 4G-4H**). Taken together, we define core human fetal and adult intestinal epithelial cell signatures that are shared *in vivo* and *in vitro,* and we demonstrate that the human and mouse fetal gene signatures differ significantly.

### Human fetal genes are not robustly activated following injury in human enteroids

We next investigated whether adult human enteroids activate a fetal transcriptional program in response to injury. Our analysis shows that adult human enteroids activate select human fetal and adult genes in an injury-specific manner (**Figure 5A-5B, Supplemental Table 8, 9**). However, we did not observe a significant enrichment (FDR q-value<0.05) of human fetal or adult genes in response to most injury conditions (**Figure 5C-5D**). Only butyrate treatment produced a significant bias towards a fetal transcriptional signature, while MA was the only condition to exhibit a significant positive enrichment for human consensus adult genes (**Figure 5E**). Enteroids treated with three conditions (5% EtOH, oxaliplatin, and 5FU) resulted in a negative enrichment for human fetal genes. Interestingly, three conditions (arsenic, DON, and ibuprofen) exhibited a negative enrichment for both human fetal and adult gene signatures. GSEA results using gene lists from only human enteroid (1002 adult genes, 761 fetal genes; **Supplemental Table 4, 5**) or primary tissue (737 adult genes, 845 fetal genes; **Supplemental Table 6, 7**) datasets did not result in significant enrichment (FDR q-value<0.05) of human fetal genes following injury, except butyrate treatment (**Supplemental Figure 5A, 5B**). Given that fetal gut development is a dynamic process that consists of multiple shifts in gene expression profiles, it is possible that human fetal regeneration signatures are specific to certain developmental timepoints. To address this limitation, we subset our enteroid and primary tissue datasets by trimester and defined consensus fetal/adult genes as described for **Figure 4** (**Supplemental Table 10**). Only butyrate treatment induced a human fetal signature when looking across trimester 1 (**Supplemental Figure 5C**) and trimester 2 (**Supplemental Figure 5D**) gene lists. Interestingly, we did observe a statistical enrichment (FDR q-value<0.05) of mouse fetal genes^16^ in human enteroids following 3/8 (Oxaliplatin, MA, 5% EtOH) treatment conditions (**Figure 5F-5G**). These results support that the human intestinal epithelium does not robustly activate human fetal transcriptional programs, but instead activates genes associated with the fetal mouse intestine for some conditions.

**Figure 5:**
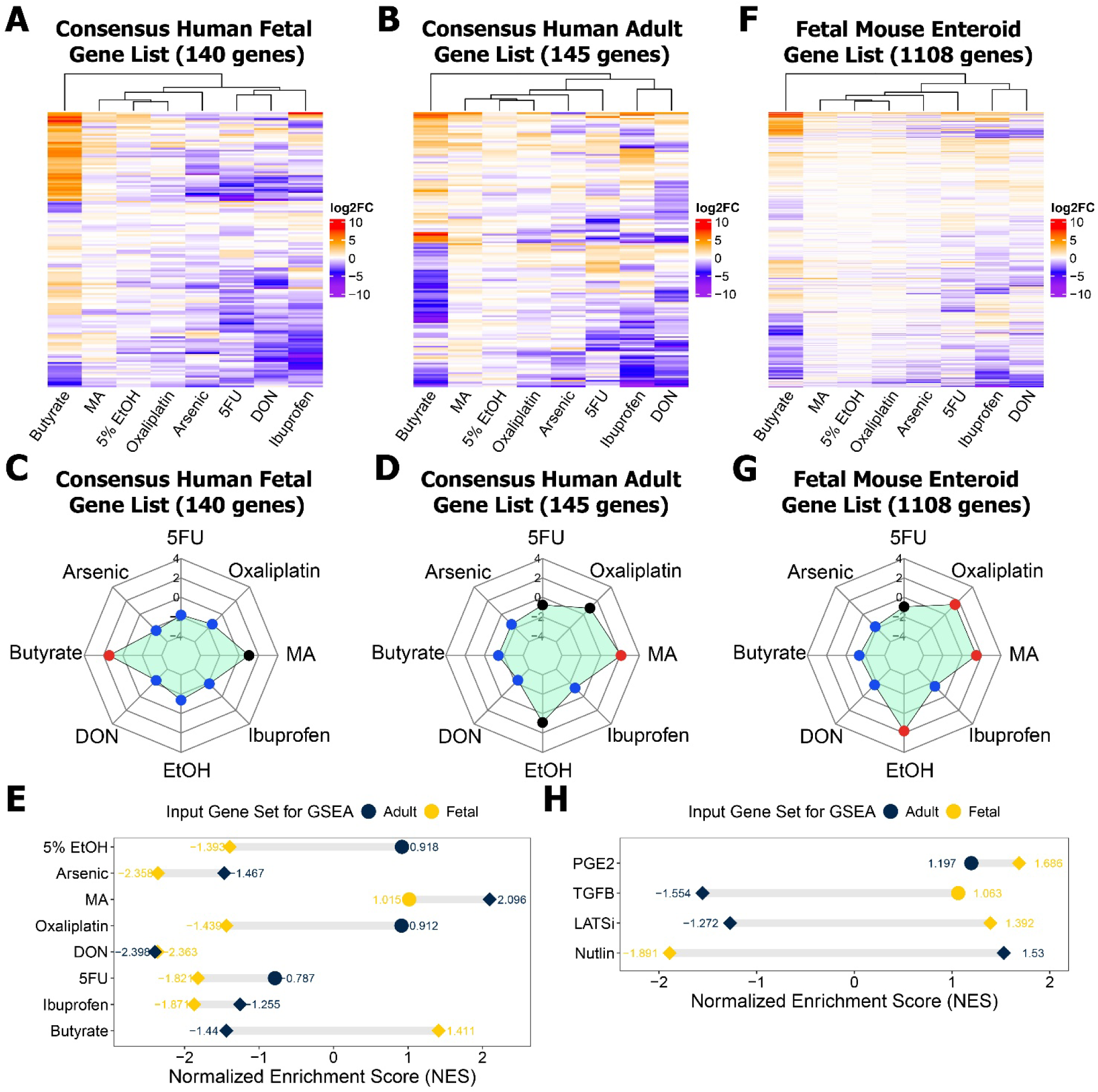
Butyrate is the only injury condition to induce a human fetal-like transcriptional signature. (**A, B**) Heatmap depicting changes in human fetal (A) or adult (B) genes in enteroids treated with eight damage stimuli. Color scale represents the log2 fold change calculated by DESeq2. (**C, D**) Radar plots visualizing NES values from GSEA analysis evaluating enrichment for consensus fetal (C) or adult (D) genes in human enteroids treated with different damaging agents. Node coloration signifies statistical significance for positive (NES > 0, FDR q-value <0.05; red), negative (NES < 0, FDR q-value < 0.05; blue), or no (FDR q-value > 0.05; black) enrichment. (**E**) Dumbbell plot visualizing results from GSEA analysis evaluating enrichment of consensus fetal (yellow) or adult (navy) genes in human enteroids treated with different injury stimuli. Datapoint shape represents whether results are statistically significant (FDR q-value < 0.05; diamond) or not (FDR q-value > 0.05; circle). (**F**) Heatmap illustrating changes in genes enriched in mouse fetal enteroids when human adult enteroids are exposed to eight injury stimuli. (**G**) Radar plots visualizing NES values from GSEA analysis evaluating enrichment for mouse fetal enteroid genes in human enteroids treated with different damaging agents. (**H**) Dumbbell plot visualizing GSEA results interrogating how activation of specific signaling pathways in human enteroids promotes a human fetal or adult transcriptional signature.

We next investigated whether activation of signaling pathways (YAP, P53, TGFβ, PGE2) induces a fetal gene signature in adult enteroids. A previous study has shown that human enteroids treated with TGFβ activate specific genes enriched in mouse fetal enteroids^17^. A gene set enrichment analysis using data from our TGFβ treated human enteroids (**Figure 3**) supports this by finding a significant enrichment of mouse fetal enteroid genes^16^ in human enteroids treated with TGFβ (**Supplemental Figure 5E**). Additional analyses found that TGFβ and YAP treatment do bias human enteroids towards a human fetal transcriptional signature (**Figure 5H**). However, activation of P53 signaling promotes an adult human transcriptional signature (**Figure 5H**). This is particularly interesting given genes downstream of P53 are most prominently activated across most injury conditions (**Figure 2C**, **Figure 3C**). Together our results show that adult human epithelial enteroids do not express human fetal genes in response to most injury stimuli and that the commonly activated P53 pathway plays a role in this by activating a more adult-like transcriptional signature.

### Arachidonic acid provides a uniquely protective effect against deoxynivalenol-induced injury

Given the broad set of transcriptional changes in response to the different injury stimuli, we wanted to determine if we could leverage data from individual damage conditions to identify injury-specific protective pathways for further exploration. To do this, we leveraged the differentially expressed gene lists for the eight injury conditions (**Figure 2A**) and performed a series of gene enrichment and pathway analyses. We observed that genes upregulated and downregulated by different injuries exhibited enrichment for biological pathway terms related to metabolism (**Figure 6A-B**). This is consistent with previous studies that have highlighted a relationship between metabolic shifts and the injury response^79–81^. Interestingly, genes downregulated by ibuprofen (p-adj = 0.037) and DON (p-adj = 0.052) were associated with arachidonic acid (AA) metabolism (**Figure 6B**). AA is an omega-6 polyunsaturated fatty acid that has been shown to promote intestinal regeneration in the mouse intestine following irradiation-induced injury^82^. Both ibuprofen and DON have been shown to affect AA metabolism under different cellular contexts. Ibuprofen is a COX inhibitor which is critical for AA processing^83,84^ while DON has been shown to reduce AA metabolites in bodily fluids^85,86^. Therefore, we hypothesized that supplementing human intestinal cells with AA will provide a unique protective effect against ibuprofen and/or DON-induced injury.

**Figure 6:**
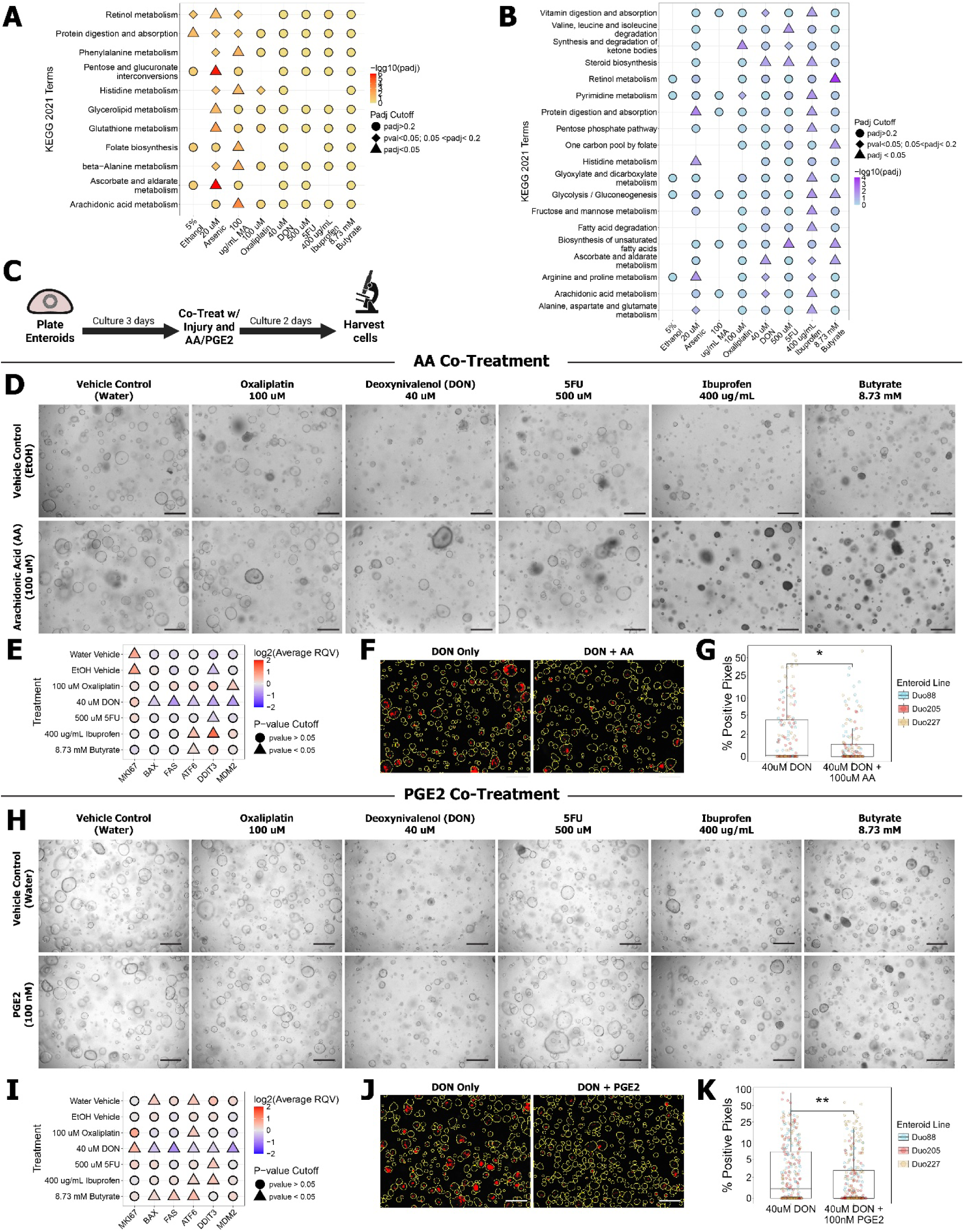
Arachidonic acid (AA) and PGE2 provide a protective effect against DON-induced injury in human enteroids. (**A, B**) Dot plot visualizing results from Enrichr KEGG pathway analysis using genes up- (A) or downregulated (B) by each treatment condition. Results have been filtered for KEGG terms affiliated with cell metabolism. Datapoint color represents the -log10(padj) and shape indicates statistical significance. (**C**) Diagram illustrating the timeline for AA and PGE2 co-treatment experiments (Created with BioRender.com). (**D, H**) Brightfield photos of human enteroids co-treated with damaging agents and AA (D) or PGE2 (H). Scale bar = 500 um. (**E, I**) Dot plot displaying differences in relative expression between injury stimuli with and without AA (E) or PGE2 (I). Coloration conveys the log2 relative change in expression that is induced by the addition of AA or PGE2 to each injury or vehicle control condition. Datapoint shape indicates statistical significance (two-sided Wilcoxon test). (**F, J**) Analyzed fluorescent images highlighting enteroid boundaries (yellow) and pixels positive for fluorescent signal (red) in damaged enteroids co-treated with AA (F) or PGE2 (J). Scale bar = 500 um. (**G, K**) Quantification of changes in the relative percentage of each enteroid that is positive for CellEvent Caspase 3/7 reagent signal when co-treated with AA (G) or PGE2 (K). Enteroid line denoted by color. Significance determined using two-sided Wilcoxon test. *p < 0.05, **p < 0.01, ***p < 0.001. All experiments were carried out in n=3 independent biological replicates, denoted as Duo88, Duo205 and Duo 227.

To test this hypothesis, we co-treated human adult enteroids with AA and several injury conditions that induced robust alterations in cell stress, apoptosis and proliferation markers (**Figure 1D, Supplemental Figure 6A**; DON, Ibuprofen, 5FU, Oxaliplatin, Butyrate). After 48 hours treatment, we isolated RNA and performed RT-qPCR for markers of cell stress (*DDIT3*, *ATF6*), apoptosis (*FAS*, *BAX*), proliferation (*MKI67*) and P53 activity (*MDM2*) (**Figure 6C-6E**). Our results show that AA treatment in the absence of injury (control groups; Water Vehicle + AA and EtOH Vehicle + AA) increased expression of the proliferation gene *MKI67* (**Figure 6E**). Co-treatment of ibuprofen and AA increased the expression of cell stress markers (*DDIT3, ATF*), which may result from an inability to process the additional fatty acid due to COX inhibition. Cells injured with DON and supplemented with AA exhibited a significant decrease in *MDM2*, cell stress, and apoptosis genes relative to cells treated with just DON (**Figure 6E**). Follow up experiments using the Caspase 3/7 detection reagent confirmed that supplementation with AA provided a unique protective effect against DON-induced injury (**Figure 6F-6G, Supplemental Figure 6C**).

PGE2 is a predominant downstream product of AA^84,87^ that functions as an important regulator of the intestine’s response to damage^40,67^. Therefore, we next sought to evaluate whether PGE2 phenocopies AA and provides a unique protective effect against DON-induced injury. To test this, we repeated the co-treatment experiments performed with AA but replaced AA with PGE2 (**Figure 6H**, **Supplemental Figure 6B**). Similar to AA, treating enteroids with DON and PGE2 uniquely decreased *MDM2*, cell stress genes, and apoptosis markers relative to cells treated with DON only (**Figure 6I**). We did not observe this same result with any of the other injury treatments tested. Subsequent experiments also confirmed that PGE2 supplementation reduced Caspase 3/7 activation by DON (**Figure 6J-6K, Supplemental Figure 6D**). These results support a model by which AA provides a unique protective effect against DON-induced injury, at least in part, through its downstream product PGE2. Taken together, these experiments highlight how our sequencing data can be utilized to identify important signaling pathways that protect intestinal epithelial cells in an injury-specific manner.

## Discussion

Recent studies, primarily performed in mice, have established that the adult intestinal epithelium can activate a fetal-like gene program in response to injury^14–20^ through a YAP-dependent process^16,17,88^. While there is robust evidence to support this model of injury-repair in a murine context, studies interrogating this phenomenon in the adult human intestinal epithelium are lacking. To fill this gap, we used adult human small intestinal organoids, termed ‘enteroids’ to identify several novel findings: 1) we characterized gene expression changes that occur in adult human enteroids treated with eight different injury stimuli; 2) we defined human fetal and adult gene signatures that are present both *in vitro* and *in vivo*; 3) we evaluated whether human fetal gene programs are activated during the initial response to injury; 4) we interrogated how different signaling pathways drive fetal-like or adult-like gene signatures. Within the experimental parameters used, we found that adult human enteroids activated a human fetal gene signature in 1/8 conditions (butyrate), whereas 3/8 injuries (5% ethanol, mycophenolic acid, and oxaliplatin) induced a fetal mouse signature^16^. Furthermore, we found that injury to the adult human epithelial cells consistently led to reduced proliferation and activation of the P53 pathway. Butyrate is a unique exception that suppresses P53 activity and promotes a human fetal gene signature.

Our results also highlight that individual injuries activate diverse gene expression programs. There were a small number of genes commonly elevated (251 genes) across 4 or more treatments, and P53 plays a prominent role in the regulation of these genes. This finding is consistent with our observation that many of the 865 genes commonly downregulated include many transcripts related to cell cycle progression and DNA synthesis, processes that have been shown to be negatively regulated by P53^61^. To directly assess the epithelial response to P53, we treated human adult enteroids with Nutlin to activate P53 and found that it promoted a more adult transcriptional signature. Interestingly, there was a positive enrichment for mouse fetal genes^16^ in human adult enteroids treated with Nutlin (**Supplemental Figure 5F**), which is consistent with previous literature that showed mouse p53 is necessary for activation of murine fetal genes^39^. These findings emphasize that P53 is a key player in how the intestinal epithelium initially responds to injury by reducing proliferation and functioning as a gatekeeper of the human fetal-adult transcriptional state. Interestingly, both YAP and TGFβ signaling have been implicated in the murine fetal transcriptional response during injury-repair^16,17,89,90^, and the current study shows that activation of YAP and TGFβ in adult human enteroids stimulates a human fetal gene signature. Together, our results suggest that a human fetal-like gene expression program can be activated in the adult epithelium under specific contexts, but this does not occur during exposure to most injury stimuli.

An important limitation of our study is that it focuses solely on how the human epithelium initially responds to injury but does not interrogate what gene expression changes occur after the injury stimulus is removed. Tissue regeneration is a temporally complex process that requires tissues to respond to damage, mount a regenerative response, and re-establish homeostasis^14^. While our study supports that human fetal genes are not robustly activated in the presence of damaging agents, it is possible that a human fetal-like gene program may be deployed at a different time during the injury-repair process, for example, once the injury stimuli are removed. Future studies may include time course studies following washout of the damaging agents to address this limitation. Additionally, previous studies have shown that communication across cell types is important for coordinating a regenerative response^15,17^. Enteroids only model the intestinal epithelium and lack additional cell populations (immune, mesenchyme, etc.) that may be important for initiating a human fetal gene signature. Future studies can address this limitation by utilizing human iPSC-derived intestinal organoid models^91^, which contain a greater diversity of cell populations, to better understand the role of cell-to-cell communication during the injury response.

An interesting finding from our study was that the mouse fetal gene list published by Yui et al.^16^ is a better indicator for the injury response than human fetal genes (1/8 conditions induced fetal human vs 3/8 conditions induced fetal mouse genes). This was particularly surprising given that we observe significant differences between genes enriched in human and mouse fetal contexts. The mouse fetal gene signature published by Yui et al.^16^ comes from E16.5 mouse enteroids (equivalent to human trimester three^92^) whereas our datasets are derived from human cells collected from trimester one and two. This observation highlights important questions for the field on how to define a “fetal” gene signature in the context of injury-repair. Are genes from specific developmental timepoints activated in the adult intestine during injury-repair (trimester 1 vs trimester 3)? Should fetal gene lists be defined using data from primary tissue or enteroids? Should a “fetal” gene be exclusive to a developmental context or simply more highly expressed in the fetal intestine? It is important to address these questions in the future to solidify our understanding of how the human intestinal epithelium responds to damage.

Finally, results from our study highlight that various injury stimuli perturb genes associated with metabolic processes in adult human enteroids. This is in concordance with prior literature that shows that metabolic processes play a critical role in how various cell types respond to damage^79,93–95^. The underlying assumption is that damaged cells undergo metabolic shifts to acquire the energy and materials needed to facilitate repair^96^. Our data is in alignment with this model by showing that DON-induced injury, which downregulates genes related to AA metabolism, can be reduced by supplementing enteroids with exogenous AA. While bulk RNA-seq played a critical role in highlighting this protective mechanism, transcriptomics is limited in its ability to characterize metabolic shifts due to its inability to capture post translational modifications, enzyme activity or metabolite abundance. Future studies can build on this work by integrating our RNA-seq datasets with metabolomics and/or proteomics data to establish a more complete perspective on how intestinal injury shapes the metabolic landscape.

Taken together, our findings provide novel insight into the molecular changes human intestinal epithelial cells undergo in response to various injury stimuli. In doing so, we created a publicly accessible database, called HD-REPAIRD (https://spence-lab.shinyapps.io/HD-REPAIRD/), that offers a rich resource for the research community to utilize our 140-bulk RNA-seq datasets generated from this study. This database includes sequencing data from 3 different patient-derived enteroid lines (2 female, 1 male) that were treated with 8 injury stimuli, 4 activators of injury-associated signaling pathways, and corresponding vehicle controls. HD-REPAIRD also provides the most robust comparison of human fetal (6 cell lines; 1 female, 5 male) and adult (4 cell lines; 2 female, 2 male) enteroids at baseline allowing users to identify human fetal enteroid genes and query for injury-induced gene expression changes all in one application. Furthermore, by defining a human fetal intestinal gene signature, we were uniquely equipped to evaluate to what degree human fetal genes are activated in adult human enteroids following injury. In doing so, our work provides significant advancement in the field by establishing that adult human enteroids do not robustly activate developmental gene programs in the presence of most injury stimuli. Ultimately, improving our understanding of the human intestinal injury response is essential for identifying damage-specific therapeutic targets and overall advancing the field of regenerative medicine.

## Availability of data and materials

The genomic datasets generated for the purpose of this study are deposited at European Molecular Biology Laboratory’s European Bioinformatics Institute (EMBL-EBI) ArrayExpress (E-MTAB-17519, https://www.ebi.ac.uk/biostudies/arrayexpress/studies/E-MTAB-17519?key=a3870b8b-48a4-4411-b081-95ebfd9ae630). DESeq2 differential expression statistics and rlog normalized counts are accessible at https://spence-lab.shinyapps.io/HD-REPAIRD/. Scripts used to analyze data and generate figures are accessible at https://github.com/jason-spence-lab/Villanueva_et_al_2026/. All other data/resources supporting this study are available upon request to the corresponding author Jason R. Spence.

## Acknowledgements

This project has been made possible in part by grants 2019-002440 (Seed Network) and 2021-237566 (Pediatric Network) from the Chan Zuckerberg Initiative DAF, an advised fund of Silicon Valley Community Foundation to J.R.S. This work was also supported in part by the Intestinal Stem Cell Consortium (U01DK103141 to J.R.S), a collaborative research project funded by the NIH National Institute of Diabetes and Digestive and Kidney Diseases (NIDDK) and National Institute of Allergy and Infectious Diseases (NIAID), and by the NIDDK (R01DK137806 to J.R.S; RC2DK140862 to J.R.S; R01DK121166 to J.R.S). J.W.V. is supported by the NIH Tissue Engineering and Regeneration Institutional Training Grant (T32-DE007057). We thank Ian Glass and the University of Washington Laboratory of Developmental Biology, who were supported by the NIH Eunice Kennedy Shriver National Institute of Child Health and Human Development (5R24HD000836). We also acknowledge the support of the University of Michigan Advanced Genomics Core for library preparation and next generation sequencing (University of Michigan, Ann Arbor, MI).

## Author Contributions

J.W.V. and J.R.S. conceived the study. J.R.S supervised research. Y-H.T and A.W. generated human enteroid lines and provided vital knowledge for this work. J.W.V., C.C., and M.B. performed enteroid maintenance and isolated RNA for RT-qPCR. J.W.V and A.V. processed samples for immunofluorescence staining and imaging. Y-H. T, A.W. and S.H. generated reagents critical for enteroid maintenance. J.W.V. completed remaining wet lab experiments and computational analyses. J.R.S. obtained funding for research. J.W.V. and J.R.S. wrote the manuscript. All authors edited, read, and approved the manuscript.

## Experimental procedures

### Human enteroid culture generation and maintenance

Human enteroid lines utilized in this manuscript were generated from de-identified primary human tissue as previously described^97,98^, and were obtained locally (Michigan Medicine), from organ donors via Gift of Life Michigan, or from the University of Washington Lab of Developmental Biology in accordance with the University of Michigan Institutional Review Board approval (IRB study number HUM00093465). To generate enteroid lines, proximal small intestine tissue from human patients was cut into pieces (approximately 0.5-1 cm length) and longitudinally cut to expose the epithelium. Tissue pieces were kept on ice and treated with dispase [StemCell Technologies, Cat # 07923] for 30 minutes, followed by 100% FBS for 15 minutes. An equal volume of Advanced DMEM/F12 [Gibco, Cat # 12634-028] and 100% FBS were added to the tissue pieces and they were subjected to vigorous pipetting to separate the epithelial layer. Epithelial structures were collected and washed with cold Advanced DMEM/F12. Cells were resuspended in Matrigel [Corning, Cat # 354230] and plated as 50 uL Matrigel droplets in 24-well plates. Enteroid maintenance media consisted of LWRN conditioned media^97,99^ combined with 2X human basal media (Advanced DMEM/F12, 2X Glutamax [Gibco, Cat # 35050-061], 2X HEPES [Gibco, Cat # 15630-080], 2X N2 supplement [Gibco, Cat # 17502-048], 2X B27 supplement [Gibco, Cat # 17504-044], 2X penicillin-streptomycin [Gibco, Cat # 15140-122], 2 mM N-acetylcysteine [Sigma, Cat # A9165-25G], 20 mM nicotinamide [Sigma, Cat # N0636-061]). Complete enteroid maintenance media consisted of 50% LWRN, 50% 2X human basal media, and 1 ng/mL EGF [R&D Systems, Cat # 236-EG-01M]. For experimental treatments in 96-well plates, enteroids were plated as 10 uL Matrigel droplets.

### Establishment of P53 reporter enteroid lines

#### Day 1

To dissociate enteroids, cells were treated with 1 mL Trypsin-EDTA (0.25%) [Gibco, Cat # 25-200-056] and incubated for 3-4 minutes at 37°C. Afterwards enteroids were broken apart by pipetting 3X and then pelleted to remove the Trypsin solution. Cells were washed using 1 mL maintenance media containing 10 uM Y-27632. Next, cells were resuspended in a viral infection solution containing 200 uL maintenance media (with 10 uM Y-27632) and 500 uL virus (∼6×10^7^ TU/mL) [BPS Bioscience, Cat #78666]. Cells were kept at 37°C for 6 hours and manually agitated every hour. After incubation, cells were pelleted and the viral infection solution was removed. Cells were resuspended in Matrigel and plated as 50 uL droplets in a 24-well plate. Cells were cultured in 1 mL maintenance media containing 10 uM Y-27632.

#### Day 2

Culturing media was replaced with fresh maintenance media containing no Y-27632.

#### Day 4

Antibiotic selection started by treating cells with 8 ug/mL puromycin [Sigma-Aldrich, P9620] in fresh maintenance media (no Y-27632).

#### Day 6

Media was changed and fresh 8 ug/mL puromycin was added (no Y-27632).

#### Day 7

Enteroid culturing media was replaced with fresh maintenance media (no puromycin or no Y-27632).

#### Day 8

Enteroids were passaged via needle sheering and plated in fresh 50 uL Matrigel droplets.

#### Day 11-Day 14

A second round of antibiotic selection was performed as described for Day 2 through Day 7.

### Luciferase assay and analysis

Following treatment, enteroids were resuspended in Advanced DMEM/F12 and added to 96-well glass bottom plates [CellVis, Cat # P96-1.5H-N] with 100 uL cell suspension per well. Brightfield images were acquired and used as input for Cellpose^42^ [v 4.0.9] to quantify the total area of the image that was occupied by enteroids. Luciferase assay was performed using the ONE-Step Luciferase Assay [BPS Bioscience, Cat # 60690-1] as described by the manufacturer’s protocol. Luminescent signal was divided by the total area occupied by enteroids to normalize for cell input.

### CellEvent Caspase-3/7 apoptosis assay and analysis

At time of injury treatment, CellEvent Caspase-3/7 Detection Reagent [Invitrogen, Cat # C10433] was added to the culturing media as described by the manufacturer’s protocol. At the end of treatment, enteroids were removed from Matrigel and resuspended in 1X PBS. Enteroids were added to 96-well glass bottom plates [CellVis, Cat # P96-1.5H-N] with 100 uL cell suspension per well. Afterwards, matched brightfield and immunofluorescence images were acquired. Brightfield images were fed into Cellpose^42^ [v 4.0.9] to identify enteroid boundaries and quantify the size of enteroids. Enteroid confidence scores were calculated based on enteroid circularity, elongation, image contrast at boundaries, and whether there were gaps in the detected enteroid. A mask was created for each brightfield image outlining the detected enteroids and this mask was overlaid onto the matched fluorescent images. For each enteroid, we counted the number of pixels that met a fluorescence intensity threshold and used this value to calculate the percentage of an enteroid that was positive for fluorescent signal. To ensure equal representation of enteroid lines when calculating significance, we randomly down sampled the enteroids used in our analyses so that there was an equal number of enteroids from each line when performing pair-wise comparisons.

### Enteroid fixation and paraffin processing

Enteroids were collected and washed with 10 mL 1X PBS. Afterwards, enteroids were pelleted and resuspended in histogel [Epredia, Cat # HG-4000-012]. Samples were kept on ice for 20-30 minutes. Enteroids encapsulated in histogel were fixed using 1 mL 10% Neutral Buffered Formalin (NBF) [Fisherbrand, Cat # 245-685] and kept at room temperature overnight. The following day, NBF was removed from samples and the enteroids were washed twice using Molecular Biology Grade Water [Corning, Cat # 46-000-CM]. Samples were dehydrated using the following alcohol gradient: 25% MeOH, 50% MeOH, 75% MeOH, 100% MeOH. Enteroids were kept in each solution for one hour and stored at 4°C in 100% MeOH until further processing. Prior to paraffin processing, dehydrated samples were transferred to 100% EtOH. Samples were then transferred into a 70% EtOH solution for submission to the University of Michigan School of Dentistry Histology Core for paraffin processing. Paraffin processing was performed using a Leica ASP300 automated tissue processor with 1 hour changes overnight. Afterwards, paraffin processed samples were embedded into paraffin blocks and sectioned to create 5um-thick sections for staining.

### Immunofluorescence staining

Slides containing enteroid sections were baked for one hour at 60°C in a dry oven. Sections were deparaffinized by washing slides twice with Histo-Clear II [National Diagnostics, Cat # HS-202] for five minutes each. Enteroids were rehydrated using an ethanol gradient consisting of two washes each for at least two minutes: 100% EtOH, 95% EtOH, 75% EtOH, 30% EtOH. Afterwards, slides were washed twice using ddH_2_O for five minutes. Slides were transferred into a 1X Sodium Citrate Buffer (100 mM trisodium citrate [Sigma, Cat # S1804], 0.5% Tween-20 [Thermo Fisher, Cat # BP337], pH=6.0) and steamed for 20-30 minutes for antigen retrieval. Immediately after steaming, slides were placed on ice to cool and washed twice using ddH_2_O. Sections were completely covered with a blocking solution (5% normal donkey serum [Sigma, Cat # D9663] diluted with a 0.1% Tween-20 [Thermo Fisher, Cat # BP337] in 1X PBS solution) and stored in a humidity chamber for one hour. Afterwards, sections were completely covered with primary antibodies diluted in blocking solution (see ‘Key Resources Table’ for details) and stored in humidity chamber at 4°C overnight. Slides were washed twice the next day using 1X PBS. Enteroid sections were covered with secondary antibody solution and incubated in humidity chamber for one hour at room temperature. Secondary antibodies were diluted in blocking solution (see ‘Key Resources Table’ for details) containing 1:1000 DAPI [Sigma, Cat # D9542]. Slides were washed three times using 1X PBS and mounted using ProLong Gold Antifade Mountant [Thermo Fisher Scientific, Cat # P369300]. Slides were stored at 4°C in the dark until imaging.

### RNA isolation and quantification

Enteroid RNA was isolated using the Norgen Total RNA Purification Kit [Norgen, Cat # SKU17270] as described by the manufacturer’s protocol. RNA concentration and purity was quantified using the Nanodrop One [Thermo Fisher, Cat# 13-400-519]. RNA integrity was quantified by the University of Michigan Advanced Genomics Core using a 4200 TapeStation system [Agilent, Cat # G2991BA].

### Reverse-transcription and quantitative PCR (RTqPCR)

Reverse-transcription (RT) was performed using the SuperScript VILO cDNA Synthesis Kit [Thermofisher Scientific, Cat # 11754250]. Quantification of gene expression by TaqMan probes was completed using the TaqMan Fast Advanced Master Mix [Thermofisher Scientific, Cat # 4444965] where values were normalized to *RPS9* (see ‘Key Resources Table’ for details). Quantification of gene expression by IDT custom primers was achieved using the QuantiTect SYBR Green Kit [Qiagen, Cat # 204145] where values were normalized to *RN18S* (see ‘Key Resources Table’ for details). Measurements were taken using a QuantStudio 7 Pro Real-Time PCR System [ThermoFisher Scientific, Cat # A43184]. Data was analyzed using our custom R package quickPCR (https://github.com/jwvillain/quickPCR).

### Bulk RNA-seq data processing and analysis

Libraries were prepared using PolyA enrichment at the University of Michigan Advanced Genomics core and subjected to 151bp paired-end sequencing using the Illumina NovaSeqXPlus system [Illumina, Catalog # 20084804]. Sequencing reads were aligned to the hg38 genome (GRCh38.v46) using STAR (v2.7.11a). Quantification was performed using Salmon (v1.9.0). Normalization and differential expression analyses were performed using DESeq2^43^ (v1.46.0) where we accounted for sequencing batch, sex, and condition in the model. For PCA visualizations, a rlog transformation was applied to the raw counts followed by batch correction for sequencing batch and sex using limma (v3.62.2). Transcripts with annotated gene names were used as input for Enrichr^58–60^ to perform gene set enrichment analyses for ‘KEGG 2021 Terms’, ‘ENCODE and ChEA Consensus TFs from ChIP-X’, and ‘GO Biological Process 2025.’ To define modules of genes with similar patterns of expression, a likelihood ratio test (LRT) was performed using DESeq2 on the raw gene count matrix. Genes with an adjusted p-value (p-adj) above 0.05 or a baseMean below 10 were removed. Genes were sorted in ascending order based on p-adj values and subset for the 3000 genes with the lowest p-adj. Clusters of genes were defined using rlog values as input for the degPatterns function (minc=50) from the DEGReport^62^ (v1.42.0) package.

### Single-cell RNA-seq data processing and analysis

Fetal (n=7) and adult (n=2) intestine single-cell datasets were generated from previously published work^56,57,100^. Fastq files were downloaded and ambient RNA was removed using CellBender (v0.3.0)^101^. Data was re-analyzed using Seurat (v5.3.0). Doublet removal completed using scDblFinder (v1.16.0). Cells with nFeature>10000, nFeature<1000, or percentage of mitochondrial reads>20% were removed. Data from remaining cells were normalized using scTransform and dimension reduction was performed using principal component analysis (PCA). Datasets were integrated using RPCA. Data was then subjected to a uniform manifold approximation and projection (UMAP) to visualize data along two dimensions. We next subset for clusters with robust expression of intestinal epithelial markers (*CDH1*, *CDX2*, and *EPCAM*) and re-clustered the data for analysis and visualization. Cell clusters were defined based on the expression of canonically expressed genes^51,56,102^. Differential expression was performed using the FindMarkers function from the Seurat package.

### Defining fetal and adult gene signatures

Human fetal enteroid genes were defined using bulk RNA-seq data where we subset for genes with a log2 fold change >1, p-adj <0.05, baseMean > 10 according to DESeq2. Human adult enteroid genes consisted of genes with a log2 fold change <-1, p-adj<0.05, and baseMean > 10 according to DESeq2. Human fetal primary intestine genes were defined using single-cell RNA-seq data. To select for cell clusters that are prominently represented in enteroids, we subset for the enterocyte, stem cell and transit amplifying cell populations. For each cluster, we filtered for genes with log2 fold change >1 and p-adj <0.05. The list of genes from each cluster were then combined into a single list. To ensure genes were lowly expressed in adult tissue, we subset for features that were expressed in <5% of all adult epithelial cells. Afterwards, we removed genes without unique ENSEMBL IDs and with log2 fold change <0 when looking at the epithelium as a whole. This same analysis was applied to identify adult primary intestine genes, except we subset for genes with log2 fold change <-1 in specified cell clusters, log 2 fold change <0 when looking at the whole epithelium, and expression in <5% of all fetal epithelial cells. Consensus fetal and adult genes were identified by selecting for genes that overlapped between the human enteroid and primary tissue gene lists.

### Broad Institute Gene Set Enrichment Analysis (GSEA)

To define lists of genes activated by injury-associated signaling pathways (P53, TGFβ, PGE2, and YAP) we filtered corresponding DESeq2 differential expression files for genes with log2 fold change >1, p-adj <0.05, baseMean > 10. The fetal mouse enteroid gene list used in this study was previously defined in Yui et al.^16^. Ranked lists from DESeq2 analyses (enteroid injury, fetal/adult enteroids, and enteroids treated to stimulate injury-associated pathways) were generated by ordering genes in descending order by the “stat” value calculated by DESeq2. Ranked lists from single-cell analyses (fetal/adult primary tissue) were generated by calculating the -log10(padj) for each gene and multiplying by the sign of fold change (+1 or -1) to determine if the gene was up- (+1) or downregulated (−1). Genes were then ordered in descending order according to this calculated metric. To avoid applying -log10 to genes with a reported padj=0, we added 1×10^-300^ to all padj values. All query and ranked gene names were standardized to ENSEMBL IDs. Query gene lists and ranked gene lists were processed by the Broad Institute GSEA tool (v4.4.0 build 18)^68,69^ using the following settings under the “Run GSEAPreranked” setting: 1000 permutations, “No Collapse”, Enrichment Statistic = “weighted”, and Max Size = 2000.

### Statistics

All experiments were completed using 3 different patient-derived enteroid lines unless noted in the figure legend. All bulk RNA-seq analyses were completed using DESeq2 whereas single-cell datasets were analyzed using Seurat. An adjusted p-value (padj) below 0.05 is considered statistically significant for genomic analyses. For cell culture experiments (CellEvent Caspase 3/7 assay, enteroid size, and P53 luciferase), statistical significance was calculated using a two-sided Wilcoxon test where a p-value < 0.05 was classified as statistically significant. *p < 0.05, **p < 0.01, ***p < 0.001, NS = not significant.

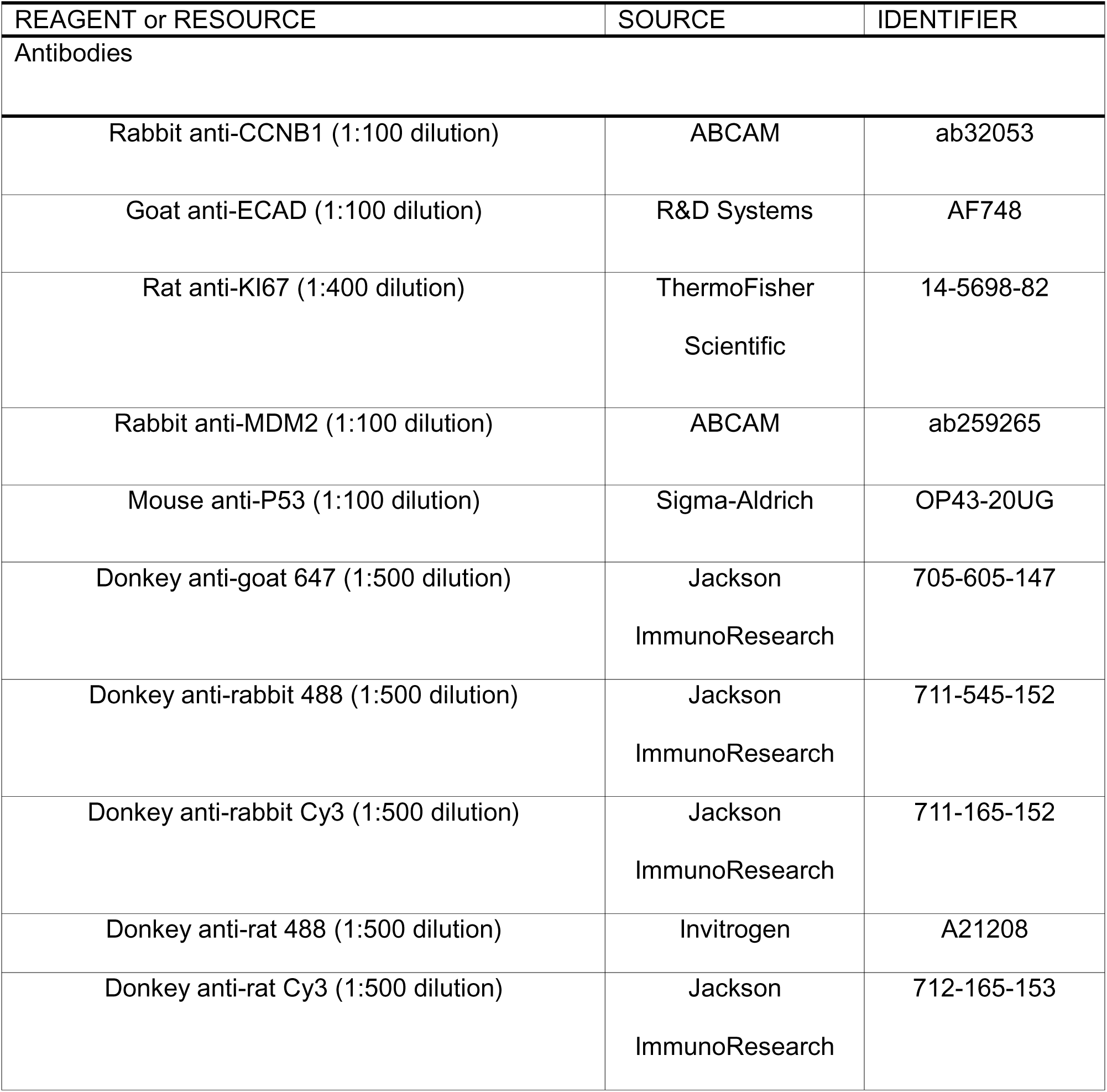
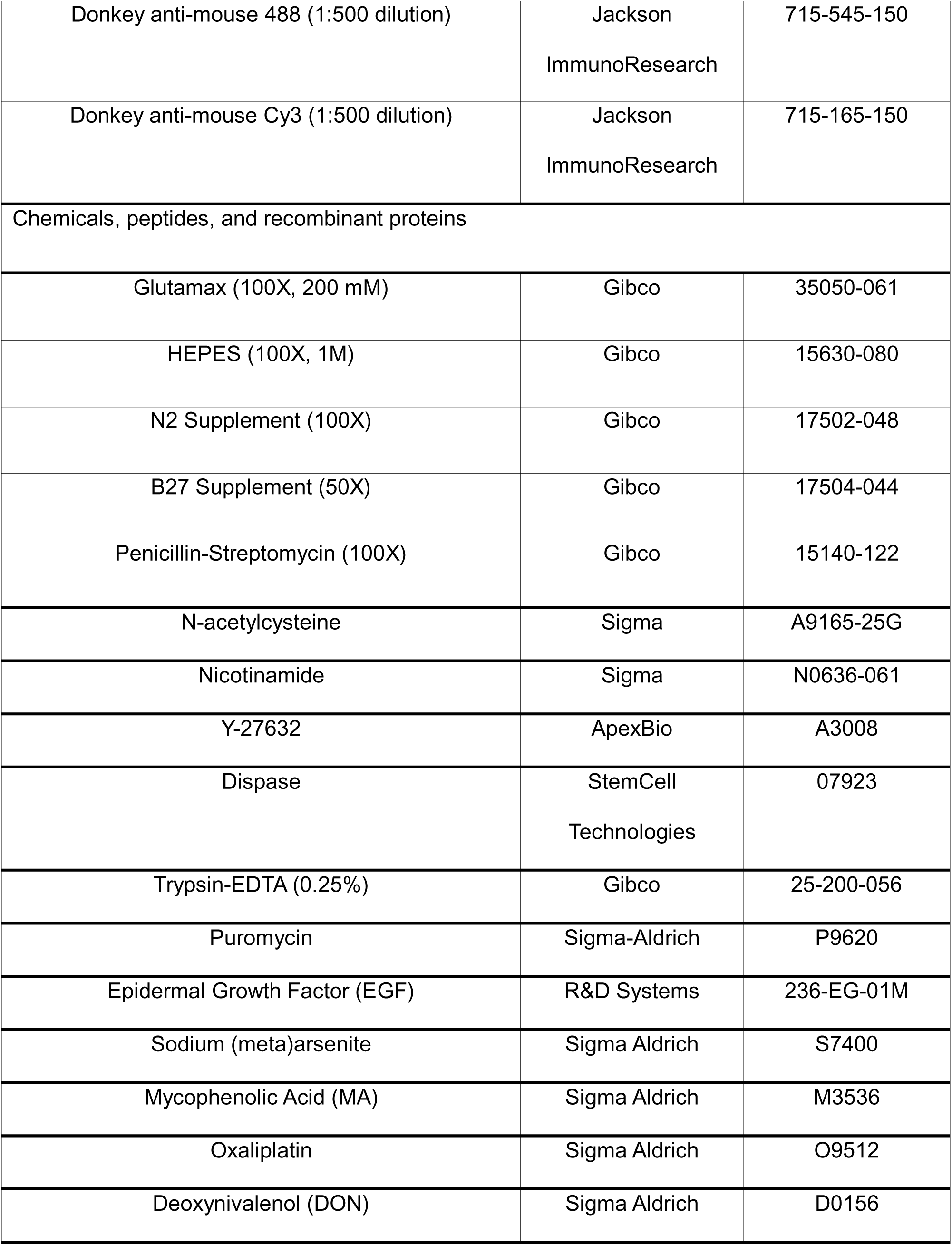
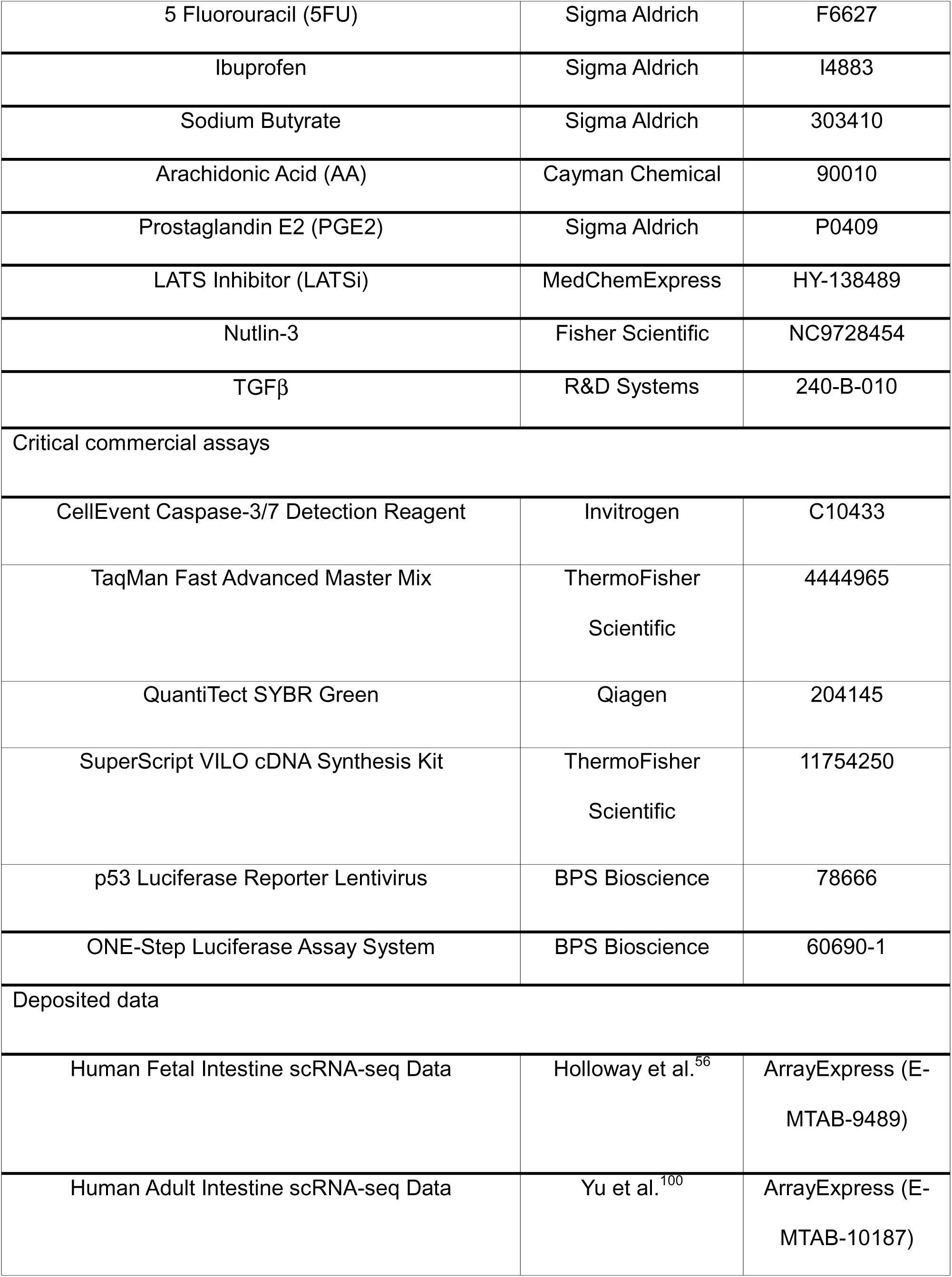
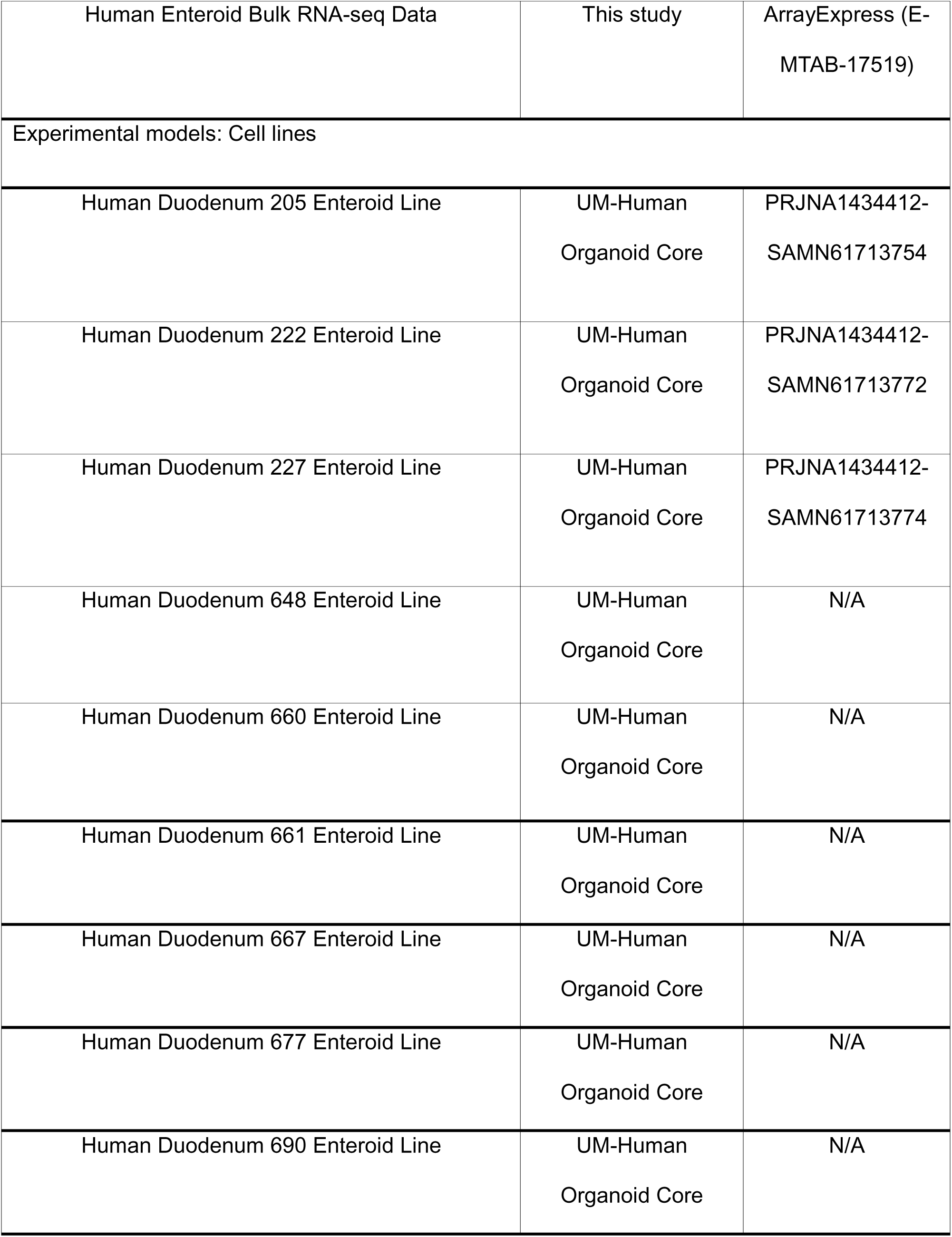
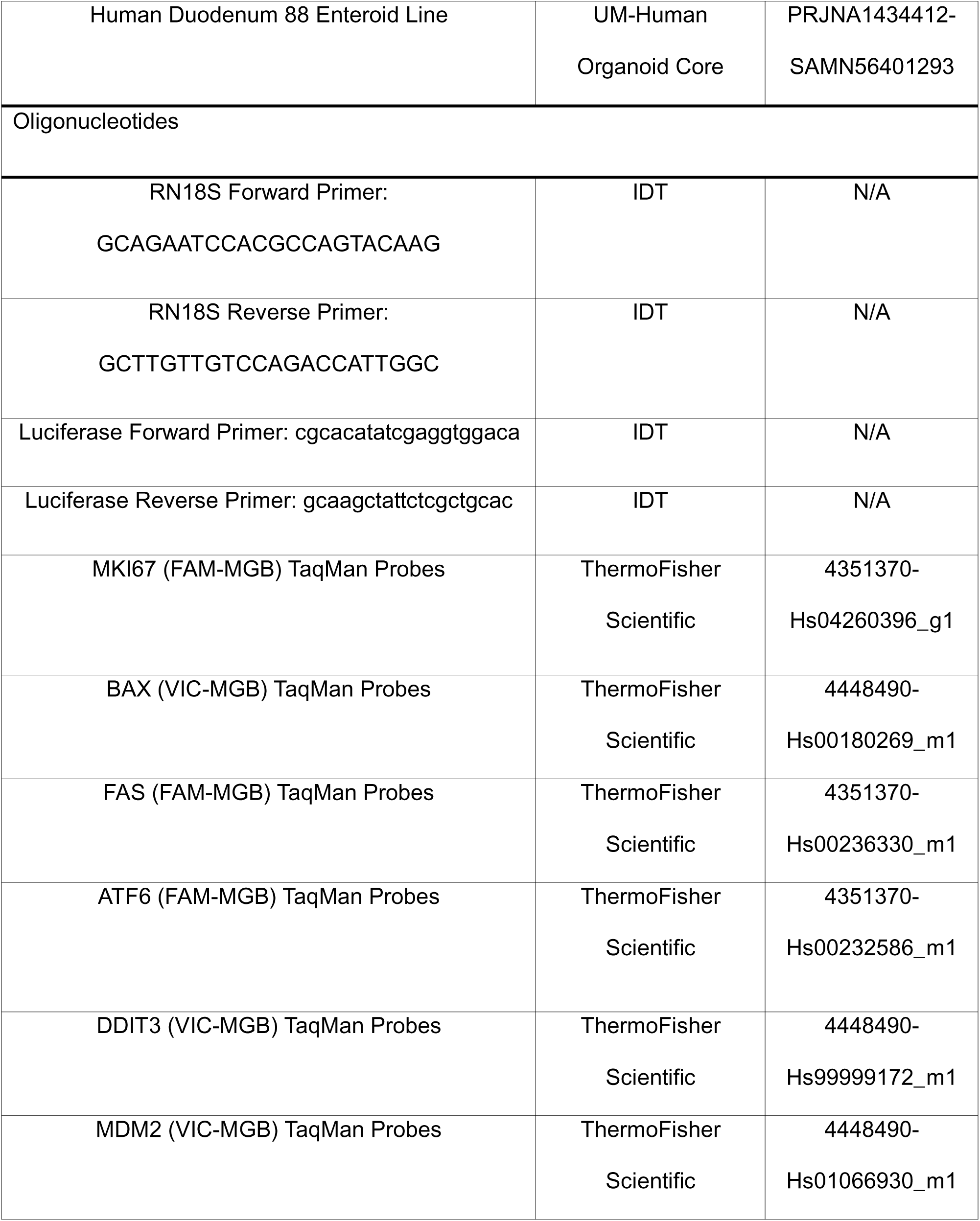
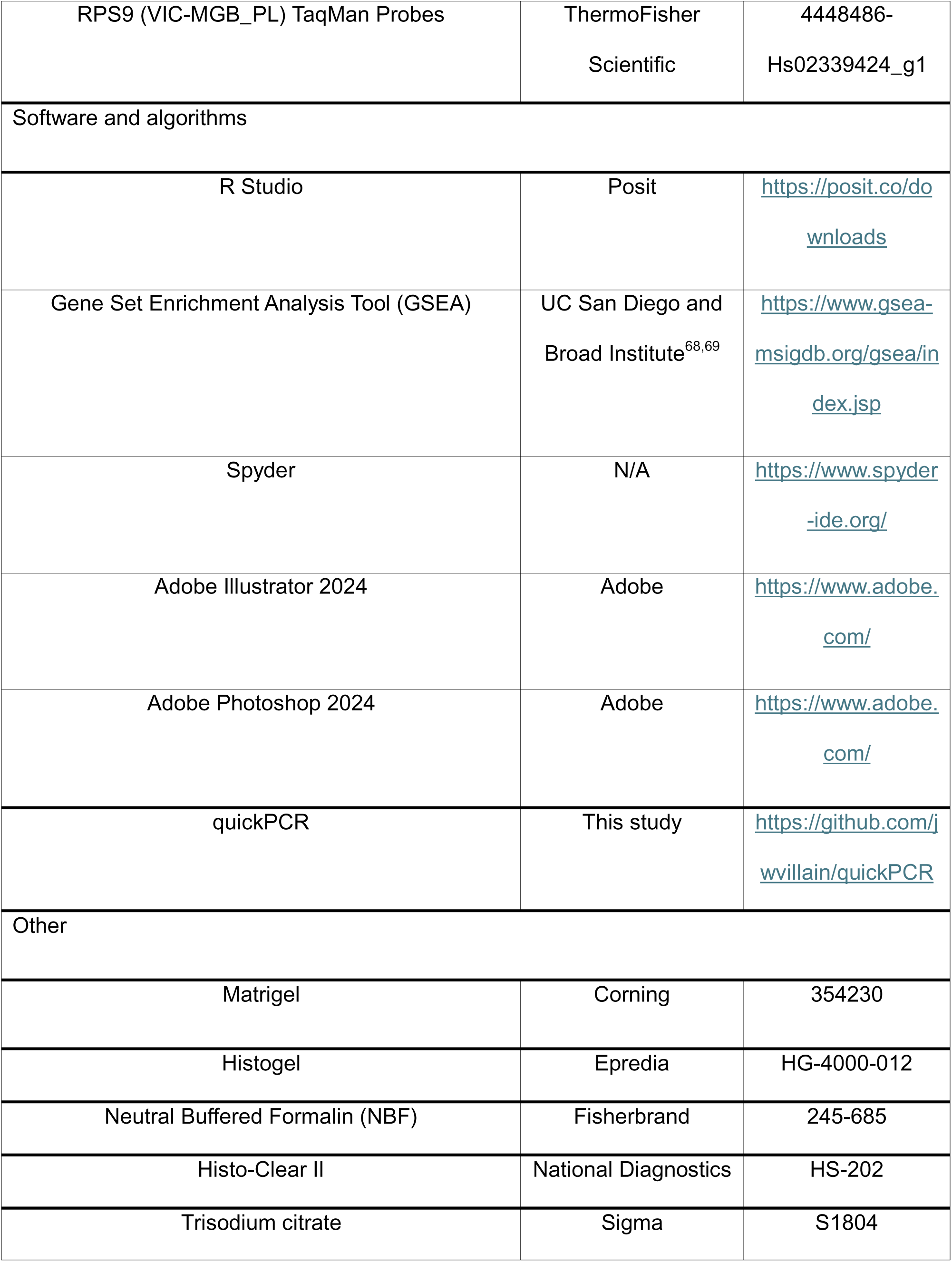
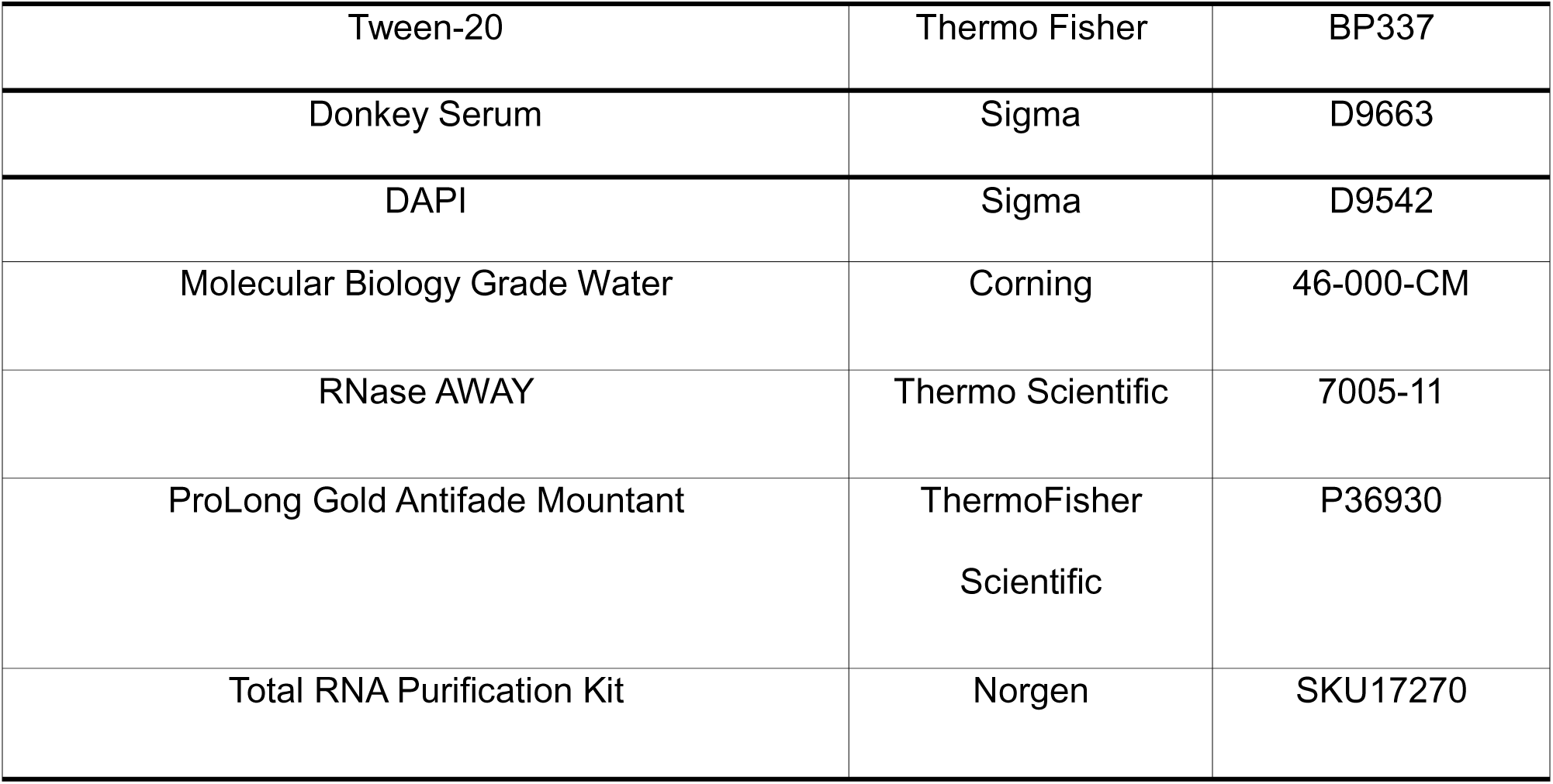
Key resources table.

