## Supplemental Figure for "Diverse Intestinal Injuries Drive Heterogeneous Transcriptional Responses and Limited Reactivation of Developmental Gene Programs in Human Enteroids"

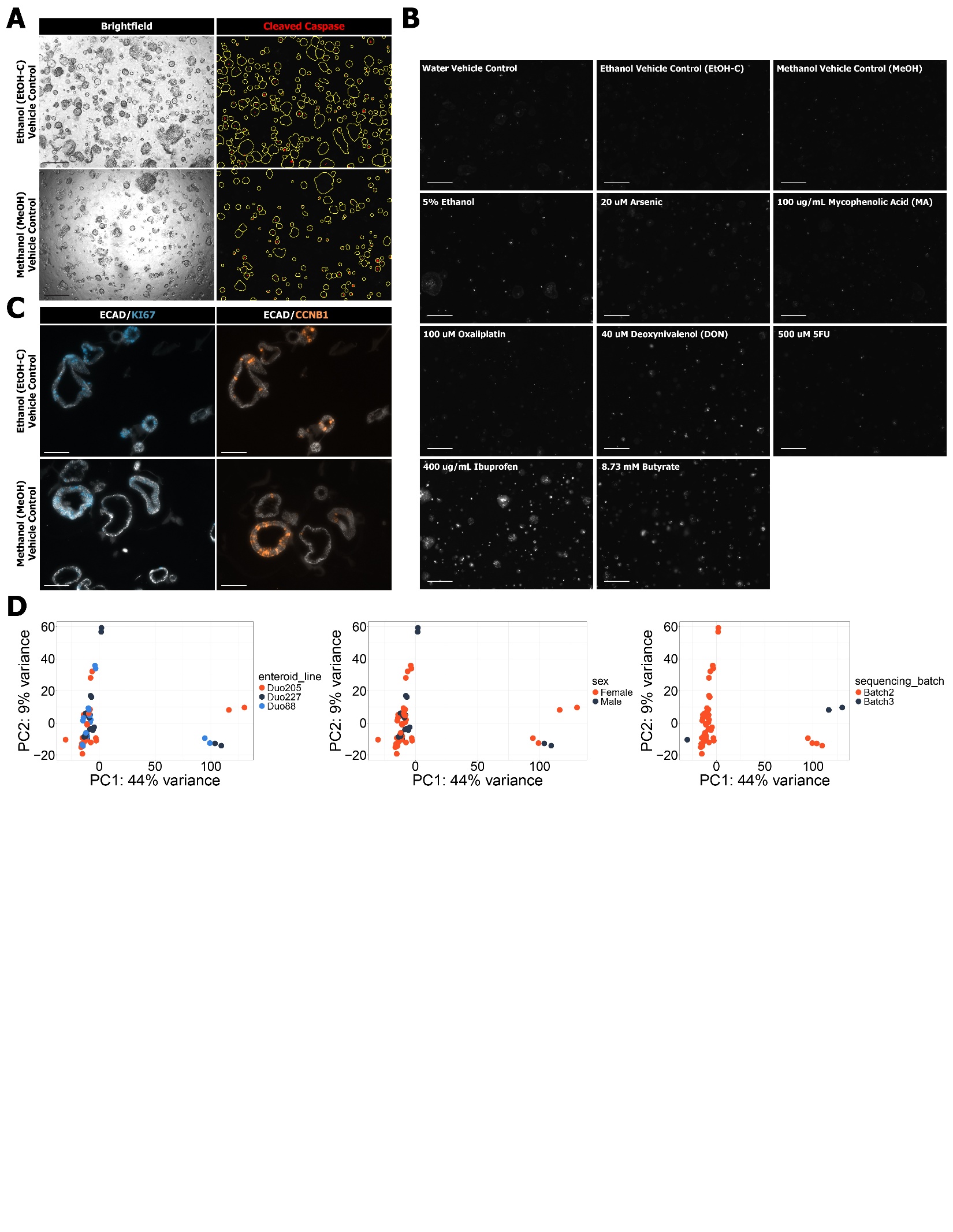

**Supplemental Figure 1**: (**A**) Brightfield (left) and analyzed fluorescent images (right; yellow = enteroid boundaries; red = pixels positive for CellEvent fluorescent signal) for human adult enteroids treated with ethanol (EtOH-C) and methanol (MeOH) vehicle controls. Scale bar = 500 um. (**B**) Raw fluorescent images of signal produced by the CellEvent Caspase 3/7 detection reagent. Scale bar = 500 um. (**C**) IF images for proliferation markers (KI67 - Blue; CCNB1 - Orange) and cell boundadies (ECAD - White). Scale bar = 100 um. (**D**) PCA generated using gene expression profiles from enteroids treated with eight injury stimuli and three vehicle controls. Datapoints colored according to sample metadata. All experiments were carried out in n=3 independent biological replicates, denoted as Duo88, Duo205 and Duo 227.

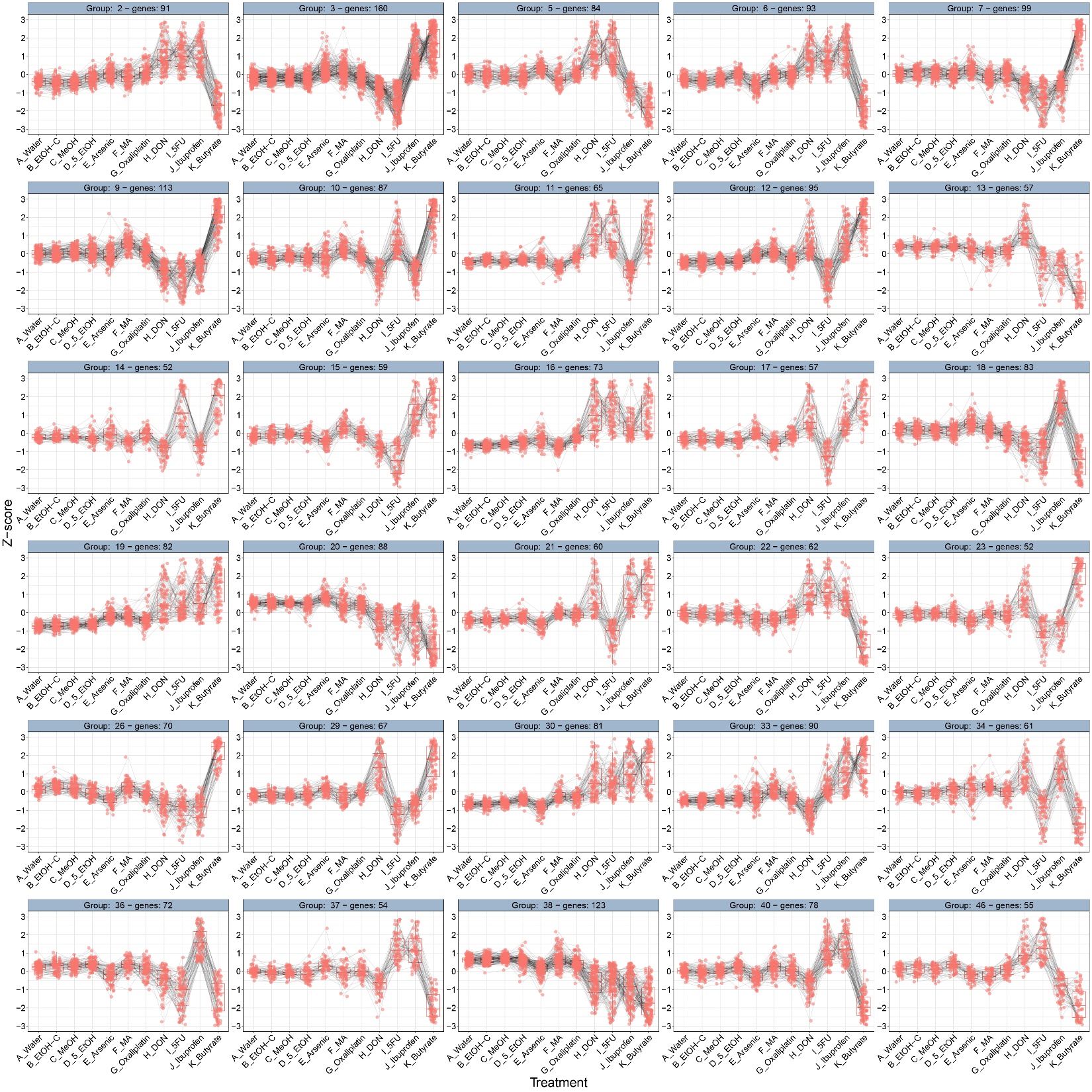

**Supplemental Figure 2**: Thirty gene modules defined by DEGReport that define clusters of genes that change in similar patterns across injury conditions. Each datapoint represents the Z-score for a gene within a given treatment condition. The black lines connects the Z-score values for the same gene across treatment conditions.

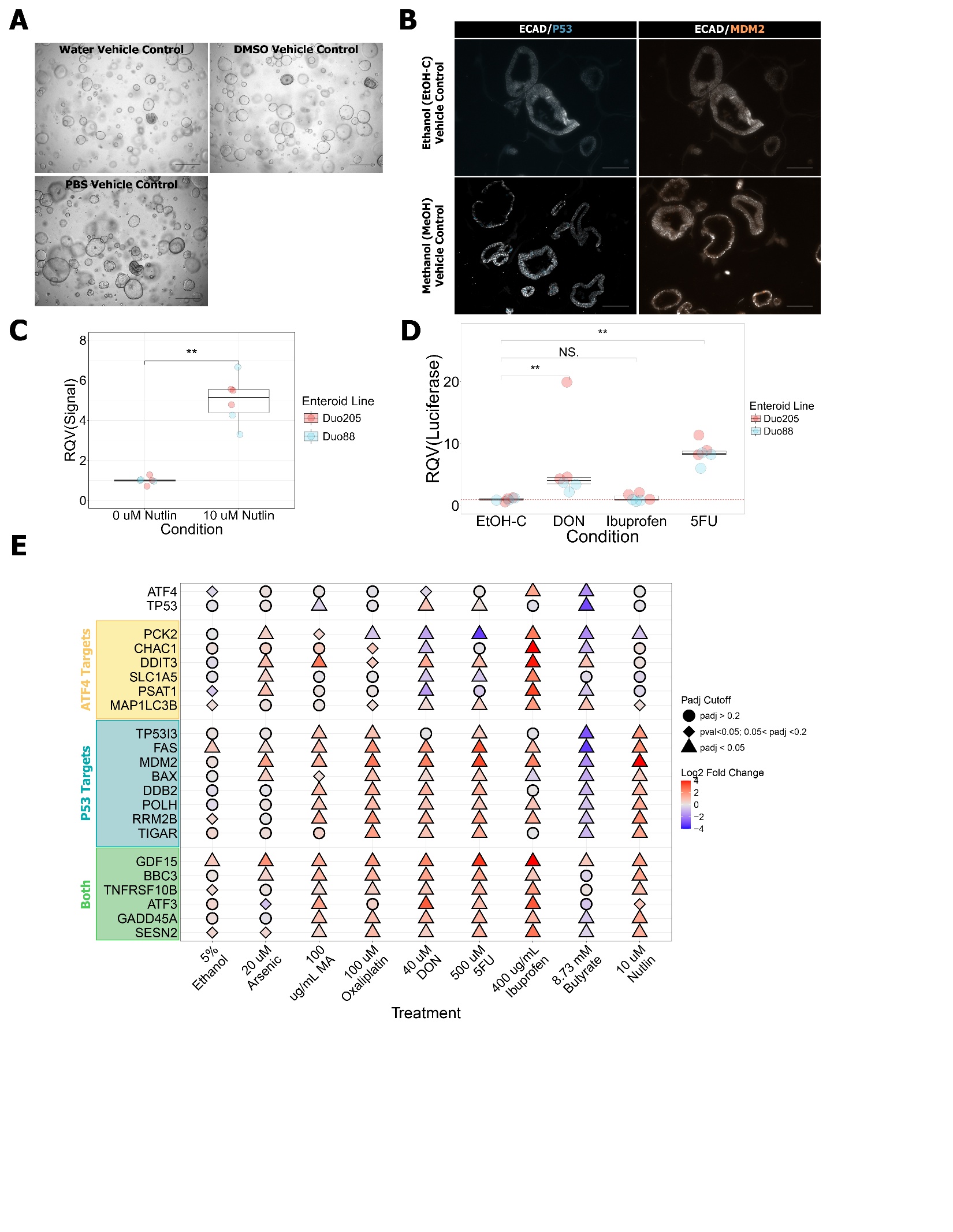

**Supplemental Figure 3**: (**A**) Brightfield images of enteroids treated with vehicle controls during experiments activating injury-associated pathways. Scale bar = 500 um. (**B**) IF images for proliferation markers (KI67 - Blue; CCNB1 - Orange) and cell boundadies (ECAD - White). Scale bar = 100 um. (**C**) Relative raw luminescent signal produced by P53 reporter lines treated with Nutlin as validation. (**D**) Relative expression of Luciferase mRNA in enteroids treated with DON, ibuprofen, 5FU, or ethanol vehicle control. (**E**) Dot plot visualizing how eight injury stimuli and Nutlin alter expression of genes activated by ATF4 (not P53), P53 (not ATF4), and both. Datapoint color represents the log2 fold change and shape indicates statistical significance. All experiments were carried out in n=3 independent biological replicates, denoted as Duo88, Duo205 and Duo 227, except for Supplemental Figure 3C and 3D with n=2 independent biological replicates.

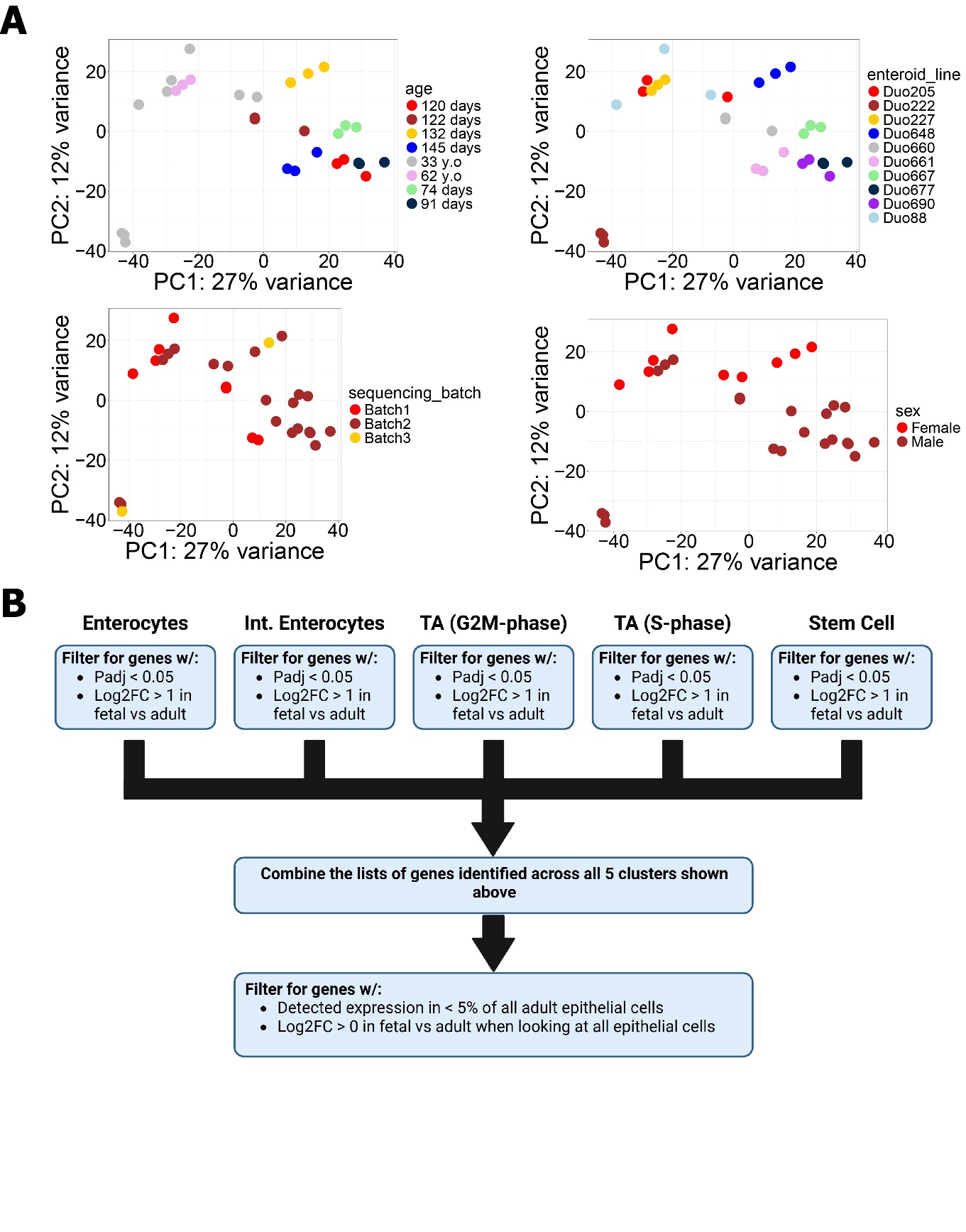

**Supplemental Figure 4**: (**A**) PCA generated using gene expression profiles from human fetal (6 lines) and adult (4 lines) enteroids at baseline (i.e. no treatment). Datapoints colored according to sample metadata. (**B**) Schematic outlining the workflow on how we identified fetal-enriched genes using scRNA-seq data from primary human tissue.

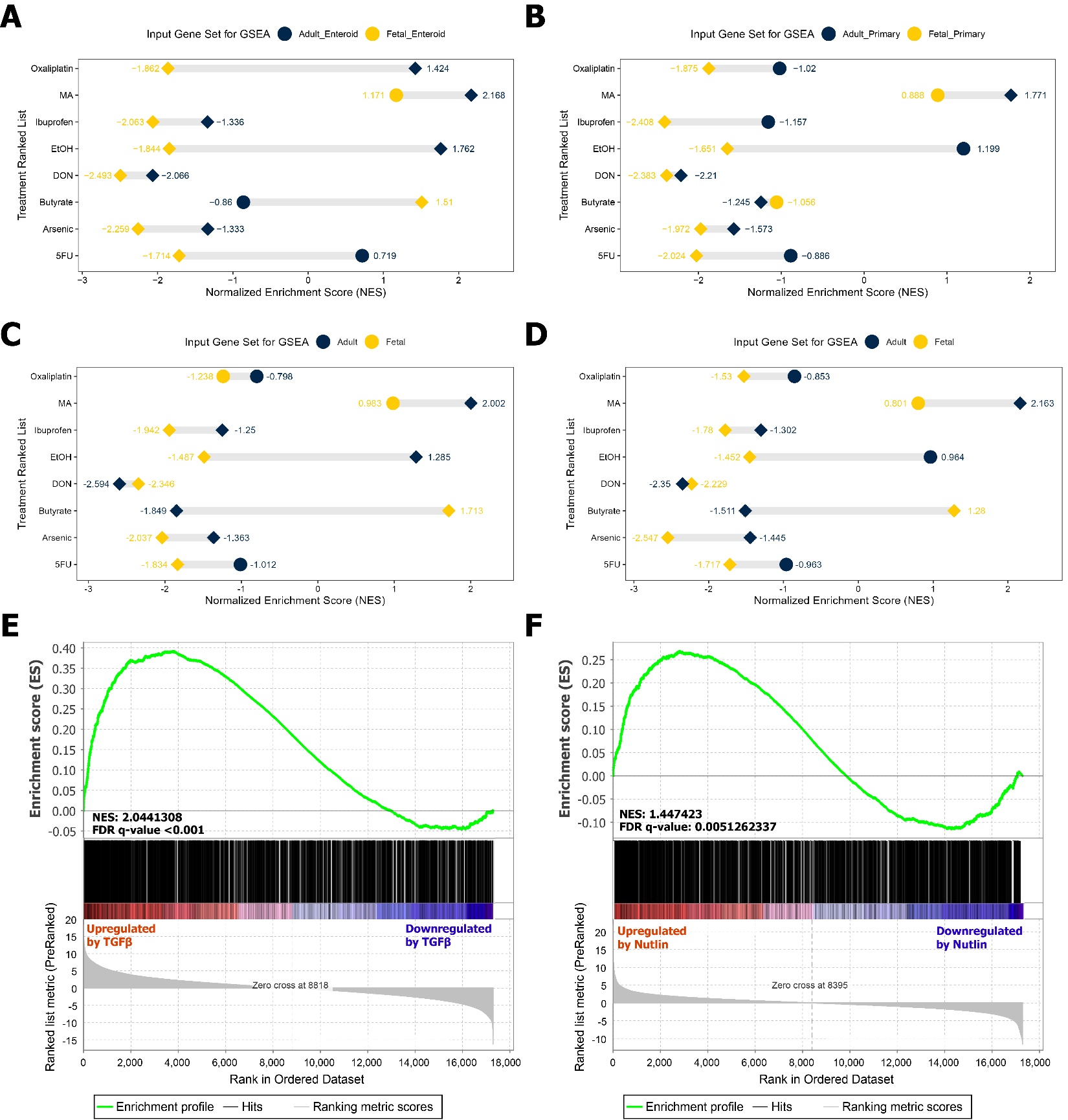

**Supplemental Figure 5**: (**A-D**) Dumbbell plots visualizing results from GSEA analysis evaluating enrichment of fetal (yellow) or adult (navy) genes in human enteroids treated with different injury stimuli. Plots utilize different criteria for defining “fetal” and “adult” as follows: Fetal and adult genes defined (A) only by the human enteroid bulk RNA-seq dataset (B) only by the human primary tissue single-cell datasets (C) by integrating enteroid and primary tissue data from trimester 1 (between 0- 97 days) (D) by integrating enteroid and primary tissue data from trimester 2 (between 98-195 days). Datapoint shape represents whether results are statistically significant (FDR q-value < 0.05; diamond) or not (FDR q-value > 0.05; circle). (**E, F**) GSEA enrichment plot evaluating enrichment for mouse fetal enteroid genes in human enteroids treated with TGFβ (E) or Nutlin (F).

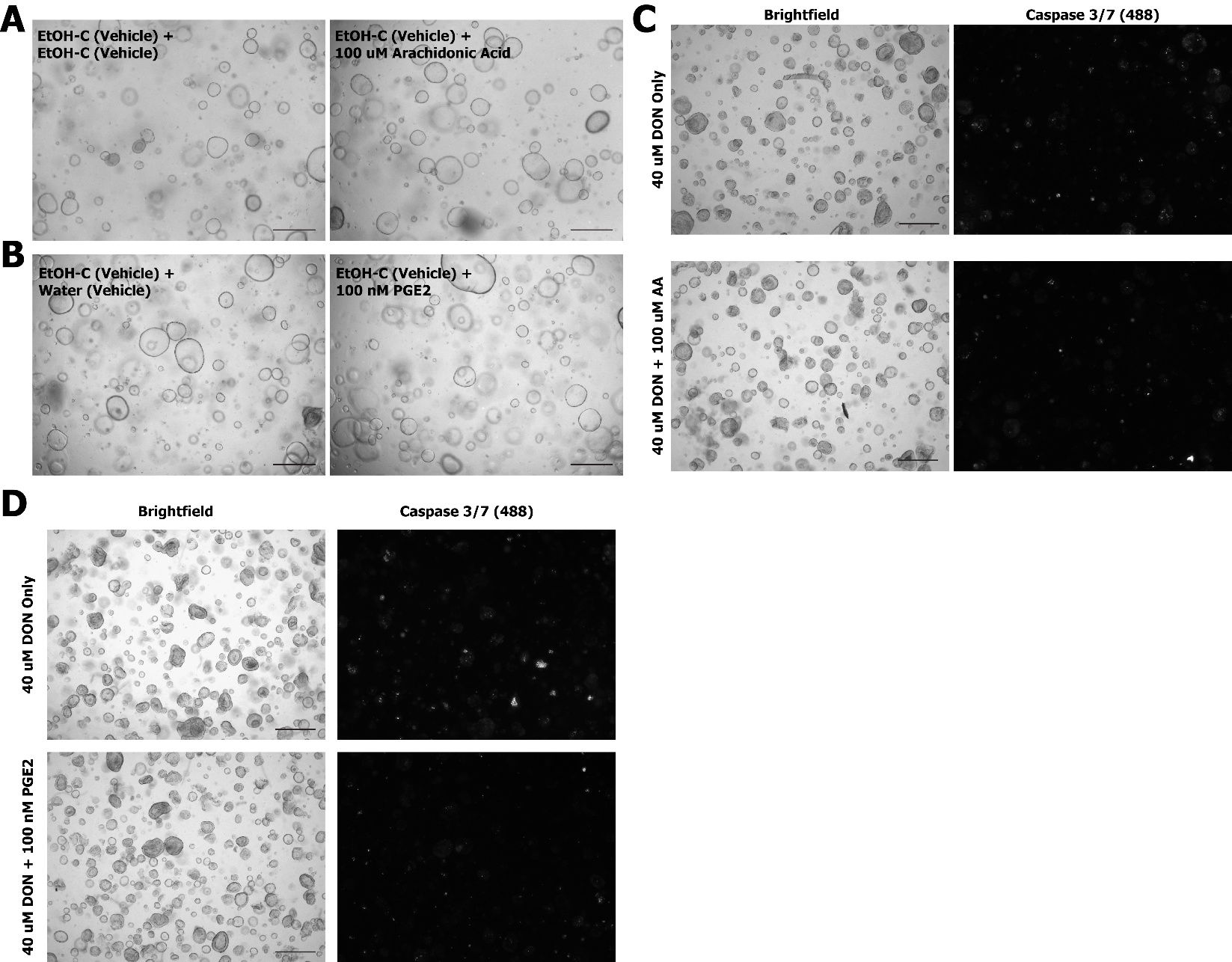

**Supplemental Figure 6**: (**A, B**) Brightfield images of enteroids treated with ethanol vehicle control during arachidonic acid (A) and PGE2 (B) co-treatment experiments. Scale bar = 500 um. (**C, D**) Brightfield (left) and CellEvent Caspase 3/7 detection reagent fluorescence (right) images for DON and arachidonic acid (C) or PGE2 (D) co-treatment experiments. Scale bar = 500 um. All experiments were carried out in n=3 independent biological replicates, denoted as Duo88, Duo205 and Duo 227.

**Supplemental Table 1**: List of 251 genes upregulated in human adult enteroids in response to 4+ injury conditions.

**Supplemental Table 2**: List of 865 genes downregulated in human adult enteroids in response to 4+ injury conditions.

**Supplemental Table 3**: Breakdown of the transcripts assigned to each of the 30 gene expression modules determined by DEGReport.

**Supplemental Table 4**: List of 761 genes enriched in human fetal enteroids relative to human adult enteroids.

**Supplemental Table 5**: List of 1002 genes enriched in human adult enteroids relative to human fetal enteroids.

**Supplemental Table 6**: List of 845 genes enriched in the primary human fetal intestinal epithelium relative to human adult intestine.

**Supplemental Table 7**: List of 737 genes enriched in the primary human adult intestinal epithelium relative to human fetal intestine.

**Supplemental Table 8**: List of 140 consensus human fetal genes that are more highly expressed in both fetal primary intestine and enteroids relative to adult.

**Supplemental Table 9**: List of 145 consensus human adult genes that are more highly expressed in both adult primary intestine and enteroids relative to fetal.

**Supplemental Table 10**: Lists of fetal and adult human intestine genes when broken down by trimester.
